# CD4 memory T cells orchestrate therapy-responsive immune niches in colorectal cancer liver metastases

**DOI:** 10.1101/2025.08.22.671704

**Authors:** Maud Mayoux, Gioana Litscher, Fabius Wiesmann, Marijne Vermeer, Pau Jorba-Adolff, Elena Guido Vinzoni, Nicolas Nunez, Colin Sparano, Viola Katharina Vetter, Ekaterina Friebel, Tobias Wertheimer, Xenia Ficht, Samira Burr, Ariana Karimi, Stanislav Dergun, Jonas Schmid, Sinduya Krishnarajah, Florian Mair, Andreas Moor, Burkhard Becher, Bettina Sobottka, Sònia Tugues

## Abstract

Colorectal cancer frequently progresses to liver metastases (CRLM), a stage with limited treatment options and poor prognosis. Neoadjuvant chemotherapy is used to control tumor growth and enable resection, yet many patients fail to respond, and the mechanisms underlying this variability remain unclear. To identify determinants of treatment response, we profiled T cell states and their spatial organization in CRLM. We found that spatial arrangement and polarization of CD4 memory T cell networks determine treatment outcome. In responders, Th1-like CD4 memory T cells organized with effector-memory CD8 T cells and antigen-presenting cells (APCs) into therapy-responsive immune niches (TRINs) that support CD4-mediated APC licensing and local immune engagement. Non-responders lacked such immune architecture, exhibiting myeloid-rich regions dominated by circulating-like CD4 memory and regulatory T cells. CD4-driven TRINs thus emerge as key determinants of chemotherapy efficacy and provide a rationale for developing biomarkers and strategies that enhance CD4-APC-CD8 crosstalk within organized immune niches.

**Statement of significance:** Th1-polarized CD4 memory T cells form therapy-responsive immune niches (TRINs) that orchestrate CD4, CD8, and APC function in colorectal cancer liver metastases, a clinically challenging and immunologically cold tumor type. TRINs define chemotherapy response and provide a mechanistic foundation for biomarker development and immunotherapy strategies designed to restore anti-tumor immunity.

## Introduction

Colorectal cancer (CRC) poses a major global health burden, with over 1.9 million new cases and 935,000 deaths annually ^1^. The vast majority of CRCs are microsatellite stable (MSS), reflecting an intact mismatch repair machinery. As a result, these tumors carry fewer mutations and respond poorly to immunotherapy, including checkpoint blockade ^2^. The principal cause of mortality in CRC is metastatic disease, most commonly to the liver, which is present in approximately 20% of patients at diagnosis and developing in up to 50% during disease progression ^3^. Because few patients present with resectable disease, neoadjuvant therapy, typically fluoropyrimidine-based combinations with oxaliplatin and/or irinotecan (FOLFOX, FOLFIRI, or FOLFOXIRI) with or without anti-VEGF or anti-EGFR agents, is used to downstage tumors and enable surgery ^3^. However, pathological responses occur in only a minority of patients ^3,4^, which is associated with improved long-term prognosis ^5–8^, underscoring the urgent need to understand why most CRLM patients fail to respond.

CD8 T cell infiltration strongly influences outcomes in CRLM ^9–11^. Both CD8 and CD4 T cells orchestrate anti-tumor immunity, with their activation and differentiation states shaping tumor progression and therapy response ^12,13^. Effector and memory CD8 T cells mediate cytotoxicity and long-term surveillance ^14–16^, while CD4 T cells enhance anti-tumor responses by licensing antigen-presenting cells (APCs), reprogramming myeloid cells, and sustaining CD8 effector and memory differentiation ^17,18^. In MSS CRC, the low neoantigen burden limits effective CD8 T cell responses and infiltration ^19–22^, an effect further exacerbated in CRLM by dysfunctional CD8 T cells and immunosuppressive myeloid cells ^23,24^. Although chemotherapy agents such as oxaliplatin and 5-fluorouracil, can induce immunogenic cell death or remodel the immune microenvironment ^25–27^, the extent to which chemotherapy efficacy is influenced by the composition, activation state, and spatial organization of CD4 and CD8 T cells in the metastatic liver microenvironment remains largely unexplored.

To address this gap, we leveraged integrated single-cell and spatial profiling to map the organization and functional states of T cells in MSS CRLM following neoadjuvant therapy. This analysis revealed that CD4 T cells dominate the lymphoid compartment of the metastatic tumor core and exhibit distinct activation and memory programs linked to therapy outcome. Patients achieving pathological response were enriched for activated Th1-like memory CD4 T cells and spatially organized therapy-responsive immune niches (TRINs), where CD4 T cells form integrated immune networks linking stem-like and effector memory CD8 T cells with specialized APCs. In contrast, non-responders lacked these coordinated structures, instead exhibiting regulatory and circulating-like CD4 memory subsets embedded within myeloid-rich, poorly interactive regions. These findings position CD4 memory T cells as central modulators of chemotherapy response in CRLM and reveal a targetable immune architecture with direct relevance for patient selection and treatment optimization.

## Results

### CD4-dominated T cell architecture defines the immune landscape of CRLM

To understand how T cell immunity shapes chemotherapy response in CRLM, we first mapped the distribution and activation states of lymphocytes across tumor regions. We performed high-throughput phenotyping on 76 matched samples obtained from the tumor center, invasive margin, and perilesional normal tissue of 30 patients with MSS CRLM. The majority had received neoadjuvant chemotherapy, with or without targeted therapy, prior to surgical resection. Histopathological response data, assessed by tumor regression grading (TRG) ^5^, were available for nearly all cases, enabling correlation between immune profiles and treatment outcomes (Fig. 1a; Extended Data Fig. 1a,b). High-dimensional flow cytometry revealed a pronounced spatial reorganization of lymphocytes in metastatic lesions. Innate-like lymphocytes (NK and γδ T cells) were largely excluded from the tumor center, along with a slight decrease in CD8 T cells and mucosal-associated invariant T (MAIT) cells (Fig. 1b; Extended Data Fig. 1c,d). In contrast, CD4 T cells and regulatory T cells (Tregs) were enriched in both the invasive margin and tumor center, where CD4 T cells accounted for nearly half of all tumor-infiltrating lymphocytes (Fig. 1b). B cells were scarce and further decreased in the tumor center (Extended Data Fig. 1e). Compared with perilesional normal tissue and the invasive margin, lymphocytes in the tumor center exhibited increased memory and tissue residency/activation features (CD69, CD45RO), but reduced effector markers (Granzyme B (GzmB), CX3CR1, KLRG1), indicating antigen experience and adaptation to a suppressive microenvironment (Fig. 1c and Extended Data Fig. 1f). The invasive margin, by contrast, was enriched in progenitor-like features, marked by high TCF1 and Eomes expression, consistent with local heterogeneity in differentiation states ^28,29^ (Extended Data Fig. 1f,g).

**Fig. 1:**
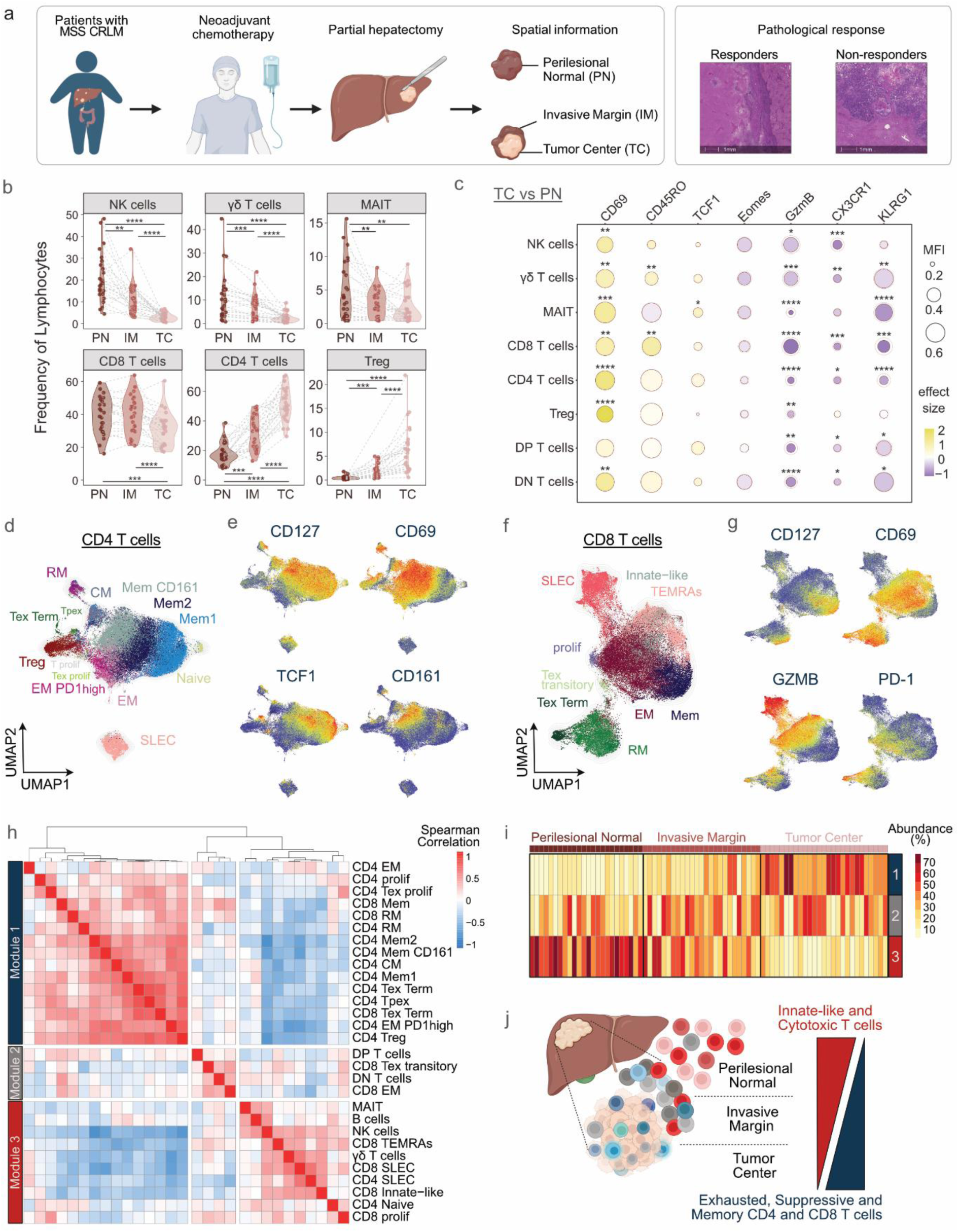
CD4-dominated T cell architecture defines the immune landscape of CRLM. (a) Overview of patient treatment history, tissue dissection into tumor center, invasive margin, and perilesional normal, and pathological response. (b) Frequencies of major lymphoid populations across regions displayed as violin plots (n = 76 matched samples from 30 patients). (c) Bubble plot showing median normalized fluorescence intensity of selected markers in tumor center (TC; black circle) versus perilesional normal tissue (PN; light red circle), colored by effect size (Cohen’s D). (d) UMAP of CD4 T cells colored by cluster. (e) UMAP of CD4 T cells colored by normalized fluorescence intensity. (f) UMAP of CD8 T cells colored by cluster. (g) UMAP of CD8 T cells colored by normalized fluorescence intensity. (h) Correlation matrix of lymphoid subset frequencies, with hierarchical clustering identifying three cellular modules (n = 76 samples from 30 patients). (i) Abundance of the three modules across perilesional normal, invasive margin and tumor center regions, each column representing an individual patient sample. Module scores represent the average normalized frequency of subsets within each module and are expressed as percentages summing to 100% per patient. See Methods for details. (j) Schematic summarizing lymphoid subsets differentially represented across the three regions. (b,c) Statistical significance was determined using paired Wilcoxon tests with Benjamini-Hochberg correction. Adjusted P value are shown as *P <0.05, **P <0.01, ***P < 0.001,****P <0.0001.

Unsupervised FlowSOM clustering further resolved this complexity, identifying diverse memory, effector, exhausted, and Treg subsets among CD4 and CD8 T cells (Fig. 1d-g). Within CD4 T cells, three predominant memory subsets were defined: Mem1 and Mem2, distinguished by distinct levels of CD69 and TCF1 expression, and a CD161^+^ CXCR6^+^ population (Mem CD161) (Fig. 1d,e and Extended Data Fig. 2a,b). The CD8 T cell compartment comprised a CD45RO^+^ CD127^+^ memory population (Mem), a CD69^+^ CD103^+^ resident memory (RM) subset, and three effector states expressing variable levels of CD69, GzmB and PD-1, including effector-memory cells (EM), short-lived effector cells (SLEC) and terminally differentiated effector cells (TEMRAs) (Fig. 1f,g and Extended Data Fig. 3a,b). We also identified distinct exhausted T cell subsets including transitory (Tex transitory) and terminally exhausted CD8 T cells (Tex Term), as well as precursor (Tpex) and proliferative exhausted CD4 T cells (Tex prolif) (Fig. 1d,f and Extended Data Figs. 2b and 3b).

To interrogate the immune organization beyond individual cell types, we analyzed the spatial co-distribution of lymphocyte subsets across tumor regions. Hierarchical clustering identified three cellular modules (Modules 1-3) (Fig. 1h), representing subsets whose abundance co-varied across samples. Module analysis and Principal Component Analysis (PCA) showed that the tumor center was dominated by memory CD4 and CD8 T cells, Tregs and exhausted T cell populations (Module 1), while cytotoxic effector and innate-like lymphocytes (Module 3) were largely excluded (Fig. 1i and Extended Data Fig. 4a-d). A complementary PCA of functional marker expression reinforced this pattern, with enriched memory and tissue-resident features and diminished effector markers across lymphoid subsets in the tumor center (Extended Data Fig. 4e,f). Terminally exhausted CD8 T cells and Tregs in the tumor center displayed elevated activation and exhaustion markers (Extended Data Fig. 4f), further supporting a functionally restrained, antigen-experienced immune microenvironment. Collectively, these findings reveal that MSS CRLM are organized around a CD4-dominated, non-cytotoxic immune landscape. The tumor center is enriched in memory and tissue-resident CD4 and CD8 T cells, Tregs and exhausted T cells, whereas the invasive margin retains more stem-like cells and the perilesional normal tissue retains predominantly effector-like lymphocytes (Fig. 1j).

### Distinct CD4 and CD8 memory T cell states define response to neoadjuvant therapy

To investigate how T cell states relate to neoadjuvant therapy, we stratified patients based on histopathological regression using tumor regression grading (TRG), a semi-quantitative measure of tumor shrinkage after therapy that correlates with long-term outcomes ^5,6^. Patients were grouped as responders (R, TRG 1 & 2), partial responders (PR, TRG 3), and non-responders (NR, TRG 4 & 5) (Extended Data Fig. 5a). We analyzed co-distribution patterns of lymphocytes within the tumor center and identified five metastatic modules (Met 1-5) (Fig. 2a), in which component subsets co-varied across samples. Responders were enriched in Met 4, characterized by a CD4 T cell memory subset (Mem2) together with CD8 memory (Mem) and effector memory (EM) T cells (Fig. 2b-d and Extended Data Fig. 5b). Non-responders, in contrast, were dominated by Met 3, featuring a distinct CD4 memory subset (Mem1) and Tregs (Fig. 2b-d and Extended Data Fig. 5b). PCA analysis of immune subset frequencies in the tumor center recapitulated these associations and further highlighted CD8 RM cells as correlated with treatment response (Fig. 2e,f). Similar patterns were observed at the invasive margin, mainly driven by CD8 T cells, whereas immune profiles of perilesional normal tissue showed no clear association with response (Extended Data Fig. 5c,d). Manual gating confirmed the strong association between CD4 Mem2 abundance in the tumor center and treatment response (R = -0.694) (Extended Data Fig. 6a-c). Notably, increases in CD4 Mem2 cells in the tumor center were accompanied by reciprocal reductions in CD4 Mem1 (Extended Data Fig. 6d), suggesting preferential activation or persistence of the CD4 Mem2 subset within the tumor core after therapy. Together, these data reveal that the therapeutic response in MSS CRLM is defined by a coordinated remodeling of the T cell compartment towards CD4 Mem2 and memory/effector CD8 subsets, and reduction of Tregs and CD4 Mem1 cells (Fig. 2g).

**Fig. 2:**
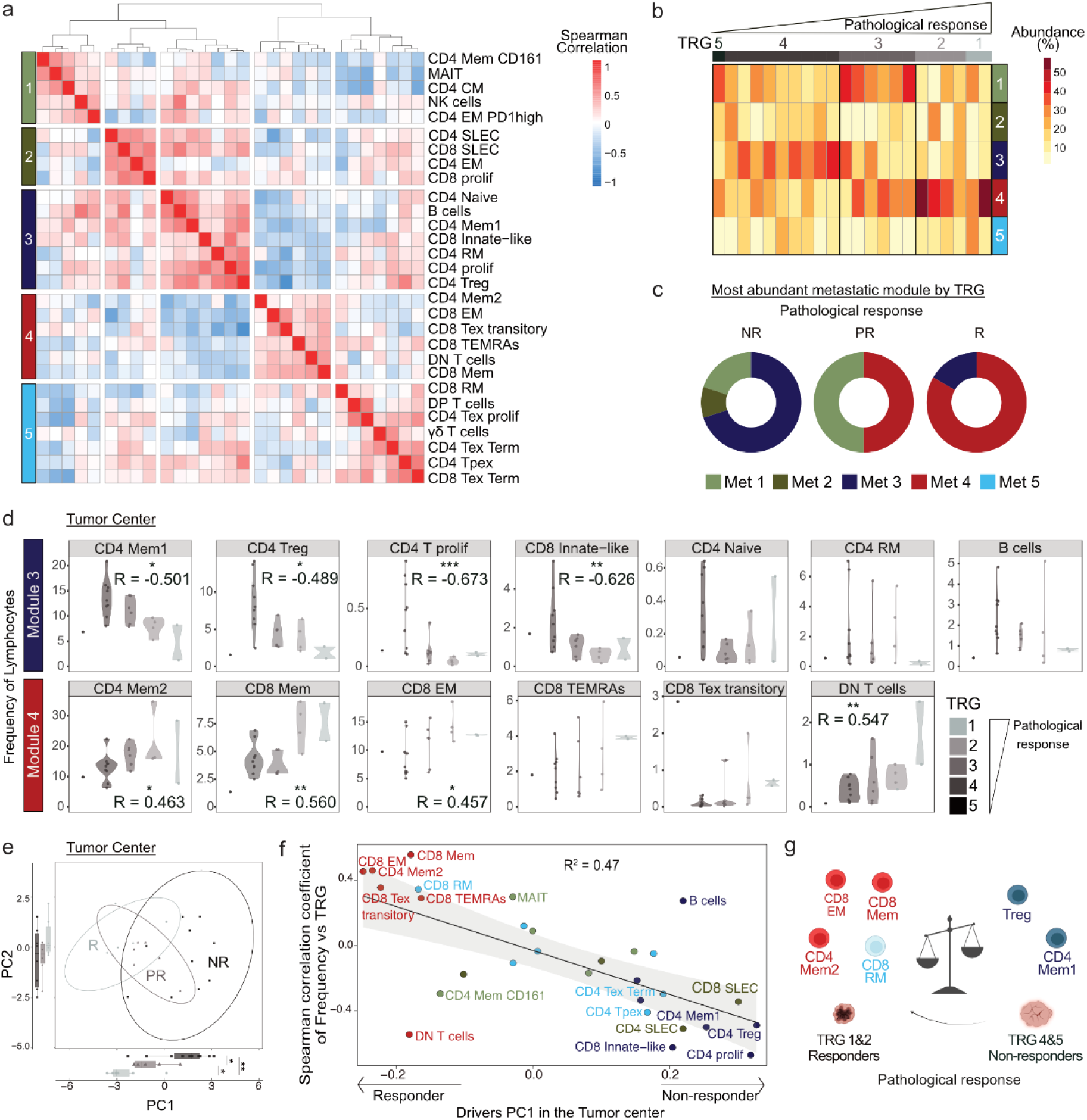
Distinct CD4 and CD8 memory T cell states define response to neoadjuvant therapy. (a) Correlation matrix of lymphoid subsets frequencies with hierarchical clustering identifying five metastatic modules in the tumor center. (b) Relative abundance of the five metastatic modules across patients, grouped by TRG score, each column represents a patient. Module scores represent the average normalized frequency of subsets within each module and are expressed as percentages summing to 100% per patient. See Methods for details. (c) Distribution of the most abundant module per patient across pathological response groups: responders (R, TRG 1&2, n=6), partial responders (PR, TRG3, n=7), and non-responders (NR, TRG 4&5, n=10). (d) Correlation between frequencies of subsets in metastatic modules 3 and 4 and TRG score. Statistical significance was determined using Spearman correlation, with coefficient (R) and exact P values shown (*P <0.05, **P <0.01, ***P < 0.001). Positive R values indicate association with low TRG (responders). (e) PCA of tumor-center immune subsets frequency per patient, grouped by response: R (TRG 1 & 2, circles), PR (TRG 3, triangles), and NR (TRG 4 & 5, squares). Boxplots display median and interquartile range of PC1 and PC2 values per group. Statistical significance was assessed by paired Wilcoxon tests with Benjamini-Hochberg correction. Adjusted P values are shown as *P <0.05, **P <0.01, ***P <0.001, ****P <0.0001. (f) Correlation between Spearman coefficient (R) of subset frequencies with TRG score (y axis) and their PC1 distance (x axis); linear regression indicates positive association (R^2^=0.47). (g) Schematic representation of lymphoid landscape differences between responders and non-responders. Analyses were performed on the tumor center samples from 22 patients with available TRG data.

**Fig. 3:**
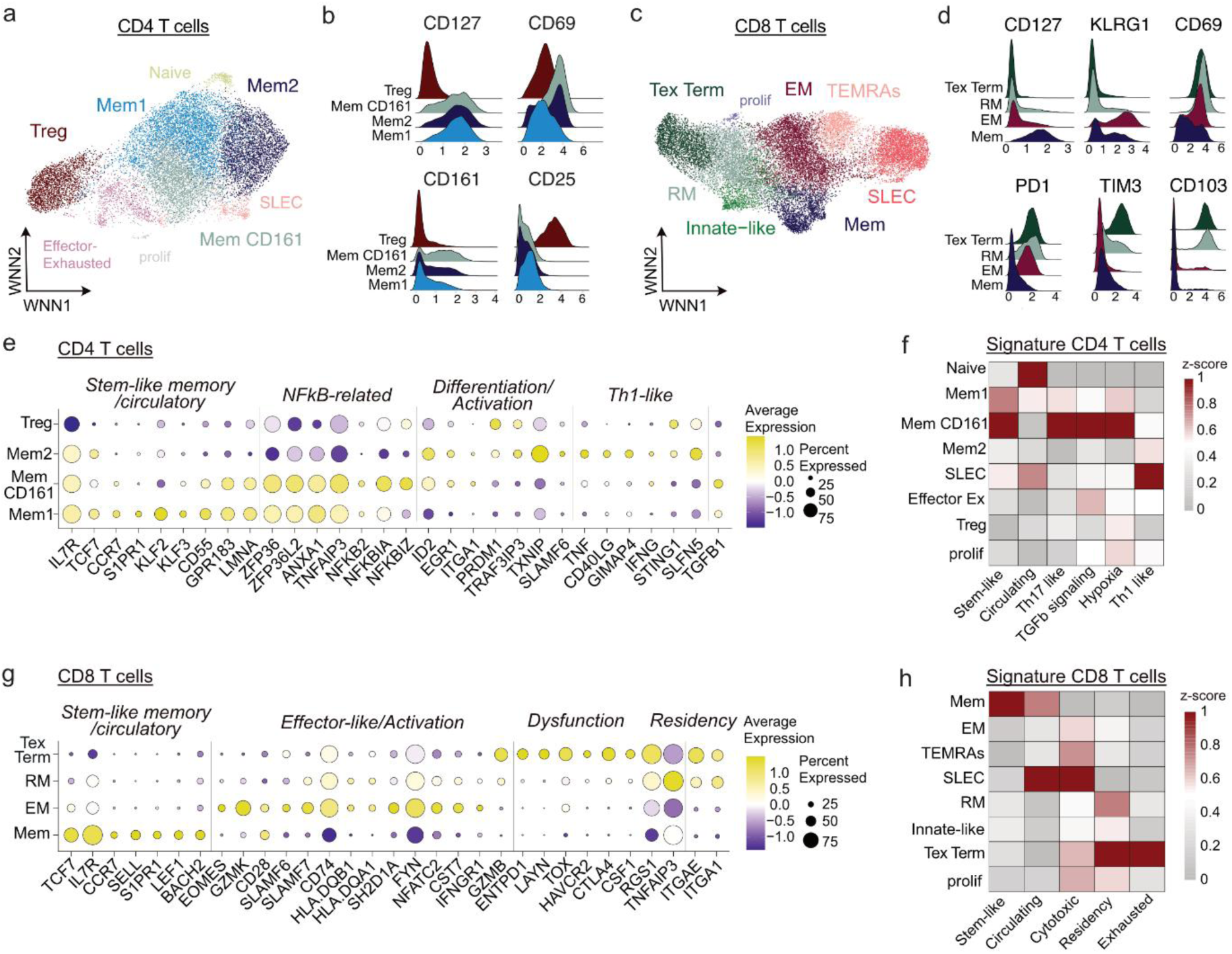
Th1-like CD4 helpers and progenitor CD8 memory programs mark therapy response in CRLM. (a) Weighted Nearest Neighbor Analysis (WNN) clustering of CD4 T cells from eight CRLM patients (15,470 cells across all regions), integrating transcriptomic and proteomic data. (b) Histogram showing z-scores of normalized protein expression (CD127, CD69, CD161 and CD25) across selected CD4 T cell subsets. (c) WNN clustering of CD8 T cells from the same cohort (19,767 cells across all regions). (d) Histogram showing z-scores of normalized protein expression (CD127, KLRG1, PD-1 and CD28) across selected CD8 T cell subsets. (e) Dotplot showing the expression of representative genes enriched in CD4 T cell subsets, based on differential expression analysis (log fold change >0.25, detection rate ≥25%). (f) Heatmap showing z-scores of predefined transcriptional signatures across CD4 T cell clusters. (g) Dotplot showing the expression of representative genes enriched in CD8 cell subsets, based on differential expression analysis (log fold change >0.25, detection rate ≥25%). (h) Heatmap showing z-scores of predefined transcriptional signatures across CD8 T cell clusters.

### Th1-like CD4 helpers and progenitor CD8 memory programs mark therapy response in CRLM

The distinct associations of T cell subsets with neoadjuvant therapy response motivated a detailed examination of the underlying molecular programs of these populations within the tumor microenvironment. We profiled eight MSS CRLM samples (six newly collected) using the BD Rhapsody platform ^30^, integrating single-cell transcriptomic, proteomic and TCR sequencing (Extended Data Fig. 7a). This multi-omic approach captured the CD4 and CD8 populations identified above (Fig. 3a-d and Extended Data Fig. 7b-d, Fig. 1), and their regional distributions mirrored the flow cytometry data (Extended Data Fig. 7e,f). Responder-enriched CD4 Mem2 cells, marked by high CD127 and CD69 and low CD161 expression (Fig. 3b), displayed a transcriptional profile of activation and Th1 polarization, including *CD40L*, *TNF* and *IFNG*-related genes (Fig. 3e,f and Extended Data Fig. 7c,g). Gene regulatory network analysis (SCENIC ^31^) confirmed activation of *STAT1*, *IRF2/3/5*, and *ETS1* regulons, reinforcing a Th1-like program (Extended Data Fig. 7h). In contrast, CD4 Mem1 cells, predominant in non-responders, exhibited a circulating, stem-like state characterized by high *TCF7* and *CCR7* expression (Fig. 3e,f), low CD69 protein (Fig. 3b), and elevated nuclear factor κB (NF-κB) and AP-1 activity (Fig. 3e and Extended Data Fig. 7h). These cells shared hypoxia- and TGF-β-associated programs with CD4 Mem CD161 cells (Fig. 3f and Extended Data Fig 7c,g), which instead exhibited a distinct Th17-like signature (Fig. 3f and Extended Data Fig. S7g) and were not linked to response (Fig. 2). Within the same compartment, Tregs were defined by high CD25 protein levels and elevated *CCR8* and *IL1R1* expression, consistent with an activated, immunosuppressive phenotype ^32,33^ (Fig. 3b and Extended Data Fig. 7c).

Parallel profiling of CD8 T cells subsets characterized the populations previously linked to therapy response, revealing a continuum of stem-like memory, effector memory and tissue resident programs. CD8 Mem T cells displayed a stem-like transcriptional program, marked by *TCF7* and *IL7R* expression, circulating features and limited clonal expansion, as assessed by TCR sequencing (Fig. 3g,h and Extended Data Fig. 7i). EM CD8 T cells expressed effector genes (*EOMES* and *GZMK*) and co-stimulatory receptors (*CD28* and *SLAMF6*), with CD69 and KLRG1 detected at the protein level (Fig. 3d,g). Their moderate *TOX* and *TCF7* expression placed them between activation and progenitor-exhausted states ^12^ (Fig. 3g). This subset showed moderate clonal expansion and shared TCRs with CD8 Mem, RM, SLEC and TEMRA subsets, suggesting multipotent differentiation potential (Fig. S7i,j). RM cells, also enriched in responders, expressed tissue residency markers (*ITGAE* and *ITGA1*) with mild dysfunction (Fig. 3g,h). In contrast, terminally exhausted (Tex Term) CD8 T cells exhibited high TIM-3, PD-1, *ENTPD1* (coding for CD39) and *TOX* expression (Fig. 3d,g), extensive clonal expansion, and TCR sharing, but were not associated with treatment response (Extended Data Fig. 7i,j and Fig. 2). Together, these findings reveal that effective anti-tumor immunity after neoadjuvant therapy is marked by Th1-like CD4 memory helpers and a spectrum of CD8 subsets spanning stem-like memory, effector memory, and tumor-reactive tissue-resident states, all engaging in coordinated local clonal expansion.

### Complementary Th1-like CD4 and effector CD8 T cell functions in therapy responders

Given the association of Th1-like CD4 Mem2 and CD8 memory/effector subsets with neoadjuvant therapy response, we next asked whether these cells differ in their functional capacity for activation and cytokine production. To test this, we performed *ex vivo* stimulation of sorted T cell subsets from the tumor center and invasive margin of 13 patients using suboptimal CD3 agonist stimulation combined with either recombinant CD40 protein or a CD28 agonist to assess helper and classical activation pathways (Fig. 4a and Extended Data Fig. 8a-d). Building on their Th1 transcriptional program, CD4 Mem2 cells, enriched in responders, exhibited Th1-type activation *ex vivo*, with higher T-bet, CD40L and PD-1 expression compared to CD4 Mem1 cells (Fig. 4b,c). They also produced more IFN-γ and TNF upon aCD3/CD40 stimulation and showed elevated production of IL-2 in response to aCD3/aCD28 (Fig. 4d), consistent with a potent helper capacity. CD8 EM cells, in turn, mirrored their transcriptional effector profile, expressing activation and cytotoxic markers (CD69, GzmK, GzmB, PD-1) (Fig. 4e,f) but showed a lower cytokine output than stem-like CD8 Mem cells (Fig. 4d). These findings reveal a functional division of labor among T cell subsets in therapy responders: Th1-like CD4 Mem2 cells provide helper activity via CD40L and cytokine signaling; CD8 Mem cells serve as a responsive reservoir, and CD8 EM cells deliver immediate effector function. This hierarchical interplay defines the mechanistic framework of effective anti-tumor immunity following neoadjuvant therapy in CRLM.

**Fig. 4:**
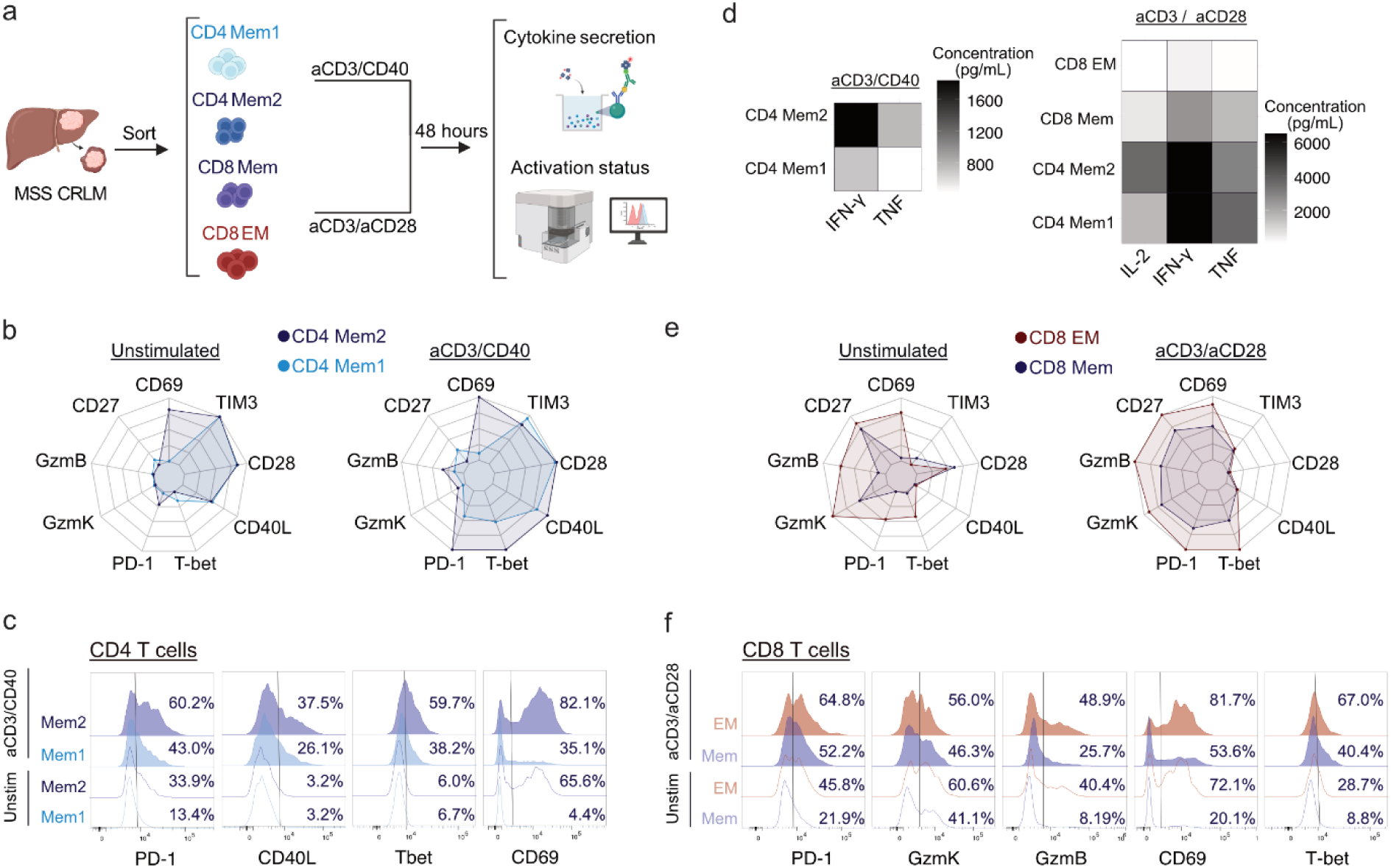
Complementary Th1-like CD4 and effector CD8 T cell functions in therapy responders. (a) Overview of the experimental design, showing sorted CD4 Mem1, CD4 Mem2, CD8 Mem and CD8 EM T cell subsets (according to the sorting strategy Extended Data Fig. 8b) and stimulated for 48 hours with suboptimal concentrations of CD3 agonist (0.5µg/mL) combined with either CD40 recombinant protein (1µg/mL; aCD3/CD40) or a CD28 agonist (1µg/mL; aCD3/aCD28). Activation status and cytokine production in the medium were assessed using flow cytometry-based measurements. (b) Radar plots showing protein expression of nine functional markers in CD4 Mem1 and CD4 Mem2 cells, unstimulated (left) or following aCD3/CD40 stimulation (right). Axes are scaled to the minimum and maximum median values across all groups and conditions. (c) Histograms showing fluorescence intensity of PD-1, CD40L, T-bet and CD69 in CD4 Mem1 and CD4 Mem2, unstimulated or following aCD3/CD40 stimulation. (d) Heatmaps showing cytokine concentration (IFN-γ, TNF and IL-2) in culture supernatant of CD4 Mem1, CD4 Mem2, CD8 Mem and CD8 EM cells after aCD3/CD40 (left) or aCD3/aCD28 stimulation (right). (e) Radar plots showing protein expression of nine functional markers in CD8 Mem and CD8 EM, unstimulated (left) or following aCD3/aCD28 stimulation (right). (f) Histograms showing fluorescence intensity for PD-1, GzmK, GzmB, CD69 and T-bet on CD8 Mem and CD8 EM, unstimulated or following aCD3/aCD28 stimulation.

### Single-cell integration reveals conserved and therapy-responsive T cell states in CRLM

To evaluate whether the T cell states identified in our cohort are conserved across patients and modulated by therapy, we integrated our Rhapsody dataset with a publicly available scRNAseq dataset from 10 treatment-naive MSS CRLM patients, comprising tumor center and perilesional tissue samples ^23^. Integration using Harmony ^34^ revealed shared CD4 T cell subsets across both cohorts (Extended Data Fig. 9a), with transcriptional signatures closely matching those defined in our dataset (Extended Data Fig. 9b, see Methods for details). In untreated patients, the tumor center was dominated by stem-like, less activated CD4 Mem1 cells, with a lower frequency of Th1-like CD4 Mem2 cells (Extended Data Fig. 9c), suggesting that neoadjuvant therapy promotes differentiation toward Th1-like helper states. In contrast, the CD8 compartment diverged substantially between untreated and treated patients, with the strongest effects seen in EM subsets (Extended Data Fig. 9d). In treatment-naive patients, data mining revealed two CD8 EM clusters (EM2 and EM3) that showed enrichment for the GzmK^+^ CD8 EM signature derived from our Rhapsody dataset (Extended Data Fig. 9e). Among these, the EM2 cluster most closely resembles the previously described GzmK⁺ CD8 EM (hC05) reported by the authors (Extended Data Fig. 9f), which represents an intermediate state bridging non-exhausted precursors and terminally exhausted CD8 T cells ^23^. In chemotherapy-treated patients, the EM2 cluster was enriched in the tumor center and invasive margin, whereas it was less frequent in untreated tumors (Extended Data Fig. 9g). These findings highlight that chemotherapy reshapes T cell immunity at multiple levels, suggesting a shift towards Th1-like CD4 states while preferentially expanding intermediate CD8 EM subsets that connect progenitors and exhausted states.

To further explore therapy-associated modulation, we examined an independent scRNAseq dataset of neoadjuvant-treated CRLM patients ^35^ stratified by treatment response. We identified a CD4 memory population (Cluster 1) expressing *IL7R*, *CD69*, *CD40LG* and low *KLRB1* (encoding CD161), resembling the CD4 Mem2 subset (Extended Data Fig. 9h,i). This cluster exhibited a Th1-like transcriptional profile and was enriched in patients with partial response (treated-PR, n=3) relative to those with progressive or stable disease (treated-PD/SD, n=6) or untreated (n=11) (Extended Data Fig. 9j,k). Together, these analyses reveal that neoadjuvant therapy reshapes conserved T cell programs in CRLM, promoting Th1-polarized CD4 helpers and modulating CD8 effector differentiation in association with clinical response.

### Spatial coordination of CD4 memory, CD8 and APCs defines therapy-responsive niches

Building on the link between Th1-like CD4 Mem2 and stem-like CD8 T cells in therapy responders, we next investigated whether their spatial organization with APCs underlies effective anti-tumor immunity ^36–39^. Using high-dimensional sequential Immunofluorescence on the Lunaphore COMET platform ^40^, we mapped immune cell distribution across tumor regions in tissue microarrays from 30 MSS CRLM patients (Fig. 5a and Extended Data Fig. 1a). Unsupervised clustering distinguished tumor cells (SATB2^+^, PanCK^+^), lymphoid populations (CD4^+^ and CD8^+^ T cells, Foxp3^+^ Tregs, CD56^+^ NK cells and CD20^+^ B cells) and diverse myeloid subsets defined by varying expression of CD68, CD11b, CD11c and HLADR (Extended Data Fig. 10a,b). Immune cells formed dense networks at the invasive margin, dominated by CD4 T cells and myeloid cells, with more modest infiltration in the tumor center (Extended Data Fig. 10c). Responders displayed higher CD4 and CD8 densities in both regions (Extended Data Fig. 10d).

**Fig. 5:**
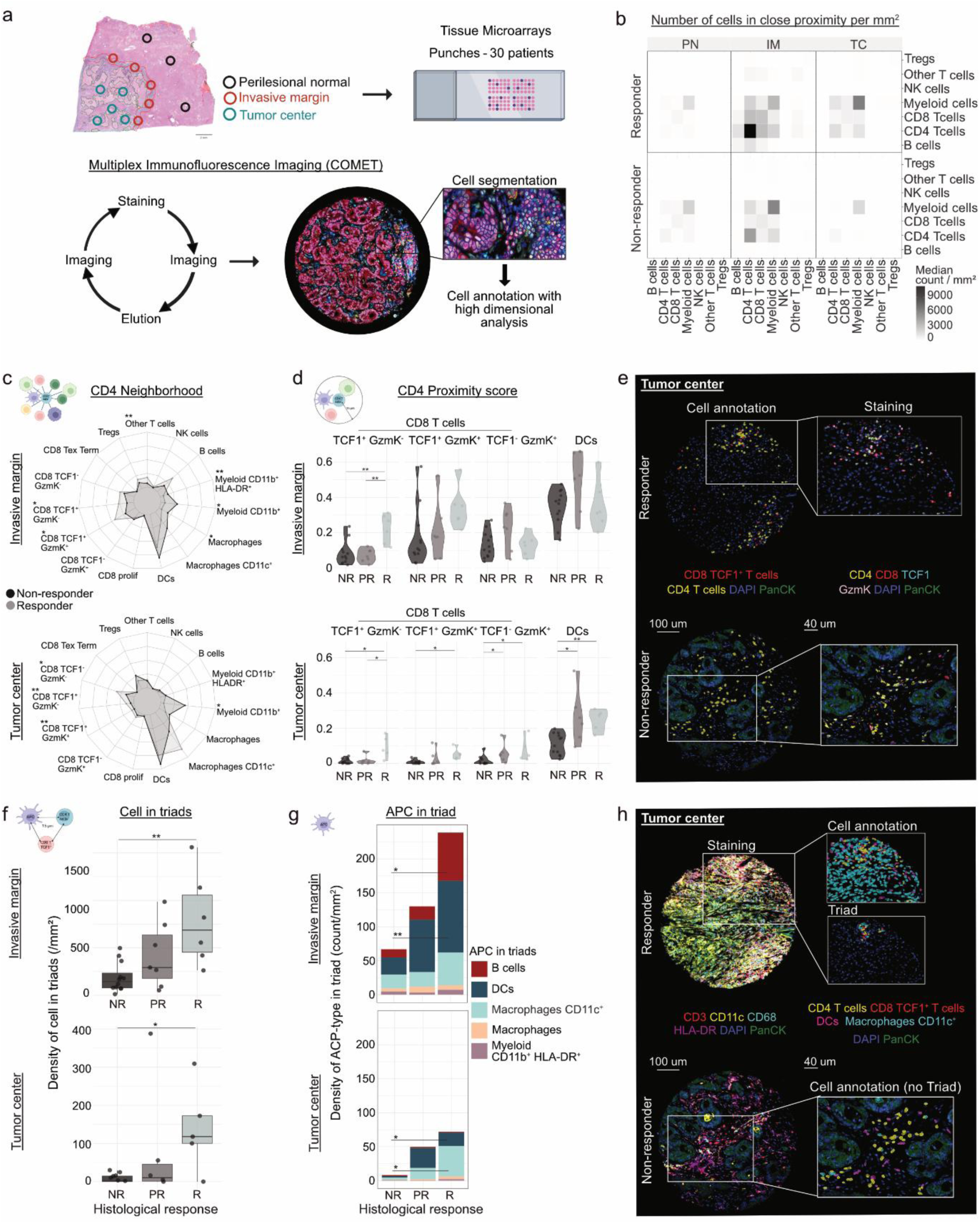
Spatial coordination of CD4 memory, CD8 T cells and APCs defines treatment-responsive niches. (a) Tissue microarrays assembled from tumor center, the invasive margin and the perilesional normal regions of 30 MSS CRLM patients (Extended Data Fig. 1a), analyzed by multiplex imaging. (b) Heatmap showing median density of cell-cell interaction (counts of proximities within a 15μm, normalized per mm²). (c) Radar plots showing mean composition of the 10 closest non-CD4 neighbors for each CD4 T cells in invasive margin and tumor center, stratified by response. Radial axes are scaled to a maximum of 35%. Statistical significance was assessed by permutation-based independence tests with Benjamini-Hochberg correction (*adjusted P < 0.05, ** < 0.01). (d) Violin plots showing proximity scores of neighbor cell types to CD4 T cells, per region and response group. Statistical significance was tested separately for each neighbor type and region, by Kruskal–Wallis followed by Benjamini-Hochberg-adjusted Wilcoxon rank-sum tests (*adjusted P <0.05, ** <0.01). (e) Representative multiplex images from tumor center of responder (TRG 2) and non-responder (TRG 4), showing CD4 T cells and CD8 TCF1^+^ GzmK^+/-^ T cells (left), with zoom-in showing selected marker staining (right). (f) Box plots showing the density (cells/mm²) of CD4 - CD8 TCF1^+^ GzmK^+/-^ - APC in triad conformation across patients, grouped by response and region. Statistical significance was assessed with unpaired Wilcoxon tests (*P < 0.05, **P < 0.01, ***P < 0.001). (g) Bar plots showing mean density of APC-type within triads across response group and region. Statistical significance was assessed with unpaired Wilcoxon tests (*P < 0.05, **P < 0.01, ***P < 0.001). (h) Representative images from tumor center of responder (TRG 2) and non-responder (TRG 4), showing full section staining (left), zoomed annotation of CD4 T cells, CD8 TCF1^+^ GzmK^+/-^ T cells, Macrophages CD11c^+^ and DCs (middle), and triad interaction (right). NR, non-responders; PR, partial responders; R, responders.

To resolve T cell states, we subclustered CD4 and CD8 T cell populations based on TCF1, CD69, and GzmK expression (Extended Data Fig. 10e-h). CD4 T cells spanned a continuum of differentiation from TCF1⁺ CD69⁻ Mem1-like cells to TCF1⁺ CD69^+^ and TCF1^-^ CD69⁺ Mem2-like populations (Extended Data Fig. 10e,f). CD8 T cells included TCF1^+^ stem-like memory (Mem) and GzmK^+^ effector memory (EM) subsets, both previously linked to therapy response (Extended Data Fig. 10g,h). Myeloid populations comprised two CD68⁺ HLA-DR⁺ macrophage subsets distinguished by CD11c levels, CD11c⁺ HLA-DR⁺ DCs lacking CD68, and CD11b⁺ HLA-DR^⁺/⁻^ myeloid cells without CD68 (Extended Data Fig. 10i,j). At the invasive margin, we observed enrichment of CD4 T cells together with TCF1⁺ stem-like and GzmK⁺ EM CD8 T cells (Extended Data Fig. 10k), suggesting active local T cell priming and differentiation. Responders exhibited higher densities of these CD8 populations and CD4 Mem2-like cells (TCF1^+^CD69^+^), along with B cells (Extended Data Fig. 10l), highlighting coordinated immune niches associated with therapy efficacy. In the tumor center, responders exhibited broader T cell infiltration (Extended Data Fig. 10l), while the region remained largely myeloid dominated, with accumulation of CD11c^+^ macrophages and CD11b^+^ myeloid cells (Extended Data Fig. 10k). Notably, CD11b^+^ myeloid cells were enriched in non-responders, whereas higher DC densities correlated with favorable outcomes (Extended Data Fig. 10l), underscoring their key role in sustaining local anti-tumor immunity ^36–39^.

Cell-cell proximity analysis revealed dense homotypic clustering of CD4 T cells and frequent close associations with myeloid populations (Fig. 5b). In responders, CD4 T cells, CD8 T cells and myeloid cells were mutually enriched in close spatial proximity (Fig. 5b), indicating the presence of organized microenvironments of immune engagement. Nearest-neighbor analysis confirmed that CD4 T cells frequently contacted DCs in both regions and CD11c⁺ macrophages in the tumor center (Fig. 5c). These neighborhoods were enriched with progenitor-like CD8 T cells (TCF1^+^ GzmK^+/-^) in responders (Fig. 5c), forming therapy-responsive immune niches (TRINs) in which CD4 T cells, CD8 T cells, and APCs interact to sustain local T cell activation. Quantitative proximity analysis using a score that corrects for cell abundance and reflects the intrinsic tendency of cells to colocalize, confirmed that TRINs exhibited significantly stronger spatial association in responders (Fig. 5d,e and Extended Data Fig. 11a). Within these niches, responders exhibited enrichment of CD4 Mem2-like cells (TCF1^+^ CD69^+^) at the invasive margin (Extended Data Fig. 11b-d) and increased CD4-DCs proximity in the tumor center (Fig. 5d), reflecting coordinated immune engagement across tumor regions.

To gain further insights into the cellular organization within TRINs, we mapped higher-order cellular triadic interactions composed of CD4 T cells, stem-like CD8 TCF1⁺ GzmK⁺/⁻ cells, and APCs (B cells, DCs, macrophages, or CD11b⁺HLA-DR^+^ myeloid cells) in close spatial proximity. These triads were markedly enriched in responders (Fig. 5f), with DC- and B cell-centered TRINs predominating at the invasive margin, and DC-CD11c⁺ macrophage-TRINs prevailing in the tumor center (Fig. 5g,h and Extended Data Fig. 11e). These findings reveal that effective anti-tumor immunity in MSS CRLM arises from spatially organized TRINs, where CD4 memory cells orchestrate local interactions with CD8 T cells and APCs. TRINs provide a structural and functional framework that sustains CD8 activation within an otherwise immunosuppressive liver microenvironment, distinguishing responders from non-responders. Thus, spatial immune organization, rather than immune cell abundance alone, emerges as a key determinant of therapeutic success in immunologically cold metastatic cancer.

## Discussion

Although chemotherapy remains the standard of care for MSS CRLM, only a minority of patients achieve meaningful tumor regression, and long-term outcomes remain poor. The limited efficacy of PD-1 blockade in these tumors ^2^ further indicates that durable antitumor immunity requires more than CD8 T cell reinvigoration alone. Here, we identify spatially coordinated immune niches enriched in Th1-like CD4 memory (Mem2) cells, stem-like CD8 T cells, and antigen-presenting cells (APCs), which emerge in therapy responders. These therapy-responsive immune niches (TRINs) integrate helper, cytotoxic, and myeloid compartments to sustain local T-cell activation and delineate the cellular and spatial architecture underlying effective anti-tumor immunity in MSS CRLM.

High-dimensional profiling revealed a spatially segregated immune landscape, with dense immune aggregates at the invasive margin and a tumor core devoid of cytotoxic lymphocytes but enriched in Tregs and terminally exhausted CD8 T cells, consistent with previous reports ^23,35^. Despite this suppressive milieu, CD4 memory T cells dominated across regions and largely retained their transcriptional programs after therapy, consistent with their relative chemotherapy resistance ^41^ and central role in coordinating immune responses and licensing APCs ^17,18,42,43^. In responders, the CD4 Mem2 subset expressed CD40L, TNF and IFN-γ, features that likely drive APC activation through CD40 signaling, enabling CD8 T cell infiltration and cytotoxic function ^44–47^. Given the well documented APC dysfunction in MSS CRLM ^19,20^, these findings underscore the importance of restoring CD4-APC crosstalk as a mean to activate myeloid and B cells. CD40 agonists, which can relieve myeloid suppression ^48,49^, induce Th1 polarization ^50^ and remodel the tumor microenvironment ^51–53^, may therefore amplify TRIN formation and enhance treatment responsiveness in otherwise cold tumors.

Spatial mapping revealed that TRINs harbor structured networks where Th1-like CD4 Mem2 cells co-localize with TCF1⁺ stem-like and GzmK⁺ effector CD8 T cells. Stem-like memory CD8 T cells showed limited clonal expansion, consistent with a self-renewing, exhaustion-resistant pool ^39,54,55^. In contrast, GzmK⁺ CD8 EM cells likely represent transitional pre-effector states ^23,55–58^ maintained in a responsive, non-exhausted state, similar to the GzmK⁺ KLRG1⁺ T cells previously linked to favorable outcomes in CRC ^35,59^. Their close spatial association with APCs suggests that TRINs function as localized activation hubs capable of maintaining effector function under low antigenic pressure ^44–47^. Given that TCF1^+^ progenitor-like CD8 T cells rely on APC-derived co-stimulation for differentiation ^60^, TRINs likely depend on CD4-mediated activation of APCs to sustain CD8 T cell renewal and function. In responders, TRINs frequently formed triadic assemblies of CD4 T cells, stem-like or GzmK^+^ CD8 T cells, and DCs in both tumor centers and invasive margins, with B cells occasionally present at the margins, potentially contributing to antigen presentation or anti-tumor IgG responses ^61^. CD11c⁺ macrophages similarly participated in TRINs in tumor cores, suggesting additional roles in antigen-presenting or chemotactic roles. Importantly, a similar triadic intratumoral architecture has been observed in mouse melanoma and hepatocellular carcinoma ^62,63^, highlighting a conserved mechanism by which CD4-APC interactions promote progenitor-like CD8 T cells differentiation and effector function.

By contrast, non-responders were enriched in Tregs and less activated, stem-like CD4 memory T cells (Mem1), associated with NF-κB activation, oxidative stress, hypoxia and TGF-β signaling, hallmarks of an immunosuppressive niche. This aligns with prior reports of SPP1^+^ macrophage-rich hypoxic niches that promote TGF-β activity ^23,24,64^ and suppress Th1 differentiation ^65^, likely limiting the emergence of Th1-like CD4 memory populations ^66,67^. Notably, tumor-specific stem-like CD4 T cells can regain Th1 polarization and enhance CD8 responses upon Treg depletion ^66^, suggesting that chemotherapy-induced immune remodeling may enable full CD4 T cell effector activation. In this context, effective CD4-mediated co-stimulation, particularly via CD40, is likely critical for APC reprogramming and sustained anti-tumor immunity. Consistent with this idea, CD4 T cells enrichment and activation of IFN-γ and TNF pathways correlate with chemotherapy responsiveness in breast cancer ^68^.

Overall, our findings position Th1-like memory CD4 T cells at the center of structured TRINs in CRLM, where they interface with progenitor-like CD8 T cells and specialized APCs to sustain anti-tumor immunity. TRINs are enriched in responders to neoadjuvant therapy and associate with favorable prognosis, highlighting the pivotal role of CD4-driven immunity in shaping treatment outcomes. By resolving the cellular architecture and signaling networks that define these niches, our work provides a mechanistic rationale for combinatorial strategies aimed at enhancing myeloid licensing and Th1 polarization to reinforce CD4-APC-CD8 crosstalk and overcome immune resistance in MSS CRLM and other immunologically cold tumors.

## Supporting information

Table 6. Differentially expressed genes in CD4 T cells from Rhapsody analysis

Table S7. Differentially expressed genes in CD8 T cells from Rhapsody analysis

Table S8. Rhapsody signature gene list

## Resource availability

### Lead contact

Further information and requests for resources and reagents should be directed to Sònia Tugues and Bettina Sobottka

### Material availability

This study did not generate new reagents.

### Data and code availability

The data generated with the Rhapsody technology have been deposite in GEO accession GSE306135 (it will be released upon publication). The code used for all analysis is available upon demand.

## Acknowledgements

We are grateful to Susanne Dettwiler and Fabiola Prutek from the Department of Pathology and Molecular Pathology (University Hospital Zurich) for their contributions to sample handling, tissue microarray production and subsequent processing. We also thank the Functional Genomics Center Zurich (University of Zurich) and the Cytometry Facility (University of Zurich) for their technical support. Specifically, we thank Qin Zhang for processing the libraries for the Rhapsody workflow and Falko Noé for pre-processing the sequencing data and sharing his expertise on computational analyses. We acknowledge Vadir Lopez-Salmeron and Eva Sofía Sánchez Quant from BD Bioscience for their valuable input in setting up the Rhapsody workflow. Additionally, we thank Victor Kreiner, Can Ulutekin Alvarez (Institute of Experimental Immunology, University of Zurich) and Florian Ingelfinger (University of Freiburg) for sharing their expertise on computational analyses.

This work was supported by grants from the Swiss National Science Foundation (PR00P3_179775 and 316030_221562) to S.T. and (320030-228252) to S.T and B.S, the Swiss Cancer Research Foundation (KFS-5420-02-2021) to S.T., the Candoc from UZH to M.M. (21-026), the Sassella Foundation to C.S. and S.T., the Vontobel Foundation to S.T., the Novartis Foundation to S.T., the Olga Mayenfisch Foundation to S.T. and the Wilhelm Sander Foundation to S.T.

## Author contributions

M.M designed, conducted, analysed the experiments and wrote the first draft of the manuscript. G.L., M.V., E.F., and F.W. supported patient sample collection and the generation of the patient bank. N.N., M.V., F.W., G.L., E.F., S.K., J.S., S.B., and C.S. assisted with flow cytometry experiments. P.J.A., E.G.V., A.K. and S.D. conducted immunofluorescence staining and imaging of slides, with P.J.A. and F.W. supporting the data analysis. T.W. assisted with the Rhapsody experiment. C.S and G.L. performed all cell sorting. V.K.V. and B.S. provided expert pathological evaluations. N.N., X.F., F.M., F.W., B.B., A.M., provided intellectual input and conceptual advice. S.T. and B.S. designed, co-supervised the study and co-wrote the manuscript with M.M., with input of all coauthors.

## Declaration of interests

The authors declare no competing interests.

## Extended data

Document 1. Fig. 1 to 11, Tables 1 to 5 and 9.

Table 6. Differentially expressed genes in CD4 T cells from Rhapsody analysis

Table 7. Differentially expressed genes in CD8 T cells from Rhapsody analysis

Table 8. Rhapsody signature gene list

## Methods

### Human specimens

All patients were enrolled at the University Hospital Zurich with pathologically diagnosed colorectal adenocarcinoma stage IV. The study was approved by the Ethics Committee (BASEC 2018-02282), and written informed consent was obtained from all participants fresh tumor and adjacent normal tissue samples from the colorectal liver metastasis surgical resection specimens were obtained by a board-certified pathologist (B.S.). Thirty colorectal cancer patients with liver metastases, including eleven females and nineteen males aged 51 to 80 years (median: 65), were analyzed using high-dimensional flow cytometry.

Patients treated with neoadjuvant chemotherapy received either Oxiplatin/Cabecetabine, Folfiri, Folfoxiri, or Folfox regiments; combined or not with targeted therapy Bevacizumab (anti-VEGF-A), Cetuximab or Panitumumab (anti-EGFR) as indicated per cohort in Extended Data Figs. 1a, 7a and 9a. Response to neoadjuvant therapy was assessed by board-certified pathologists using the TRG according to the Rubbia-Brandt classification system ^5^ based on the proportion of tumor cell and fibrosis observed in H&E stained CRLM FFPE sample. Patients were categorized as responders (TRG 1&2), partial responders (TRG3) and non-responders (TRG 4&5). The flow cytometry analysis and multiplex imaging were performed on a cohort of 30 patients (Extended Data Fig. 1a). Among the 30 patients analysed, two patients were treatment-naive, and three did not have TRG information. Thus, a total of 25 patients were included in the analysis of the relationship between response to neoadjuvant therapy and the immune landscape based on TRG. The clinical characteristics and sample regions available for each patient are summarized in Extended Data Fig. 1a. Eight colorectal cancer patients with liver metastases (Extended Data Fig. 7a) were included in the sequencing analysis using Rhapsody technology, including four females and four males, aged 36 to 70 years (median: 58).

All patients had been treated with NAC, and patients 25 and 28 were also part of the cohort of 30 patients analyzed by high-dimensional flow cytometry. The clinical characteristics and sample regions available for each patient are summarized in Extended Data Fig. 7a. Thirteen colorectal cancer patients with liver metastases (Extended Data Fig. 9a), including four females and nine males, age 41 to 77 years (median: 63), were included in the ex vivo stimulation experiment. The clinical characteristics and the sample regions available for each patient are summarized in Extended Data Fig. 8a.

### Single-cell collection

Fresh tumor and adjacent liver tissue were collected and processed shortly after surgery. The invasive margin was separated from the tumor center using a scalpel. All three regions were individually cut into small pieces and transferred into 2.5 ml of digestion medium per 0.5 g of tissue. The digestion medium consisted of RPMI-1640 medium (Thermo Fisher Scientific, #31870-025) supplemented with 10% fetal bovine serum (FBS; Thermo Fisher Scientific, #A5209402), 1 mg/ml Collagenases IV (Worthington, #LS004191), and 0.25mg/ml DNase I (Sigma-Aldrich, #DN25-5G). Tissues were enzymatically digested in a 6-well plate for 1 hour at 37°C with constant shaking at 90 rpm. The digestion was stopped with cold PBS containing 2 mM of EDTA (VWR chemicals, #E177). The digested tissue was filtered through a 100 µm cell strainer (Chemie Brunschwig, #7002109), and the resulting cell suspension was centrifuged at 300 g for 10 minutes at 4°C. The supernatant was discarded, and the pelleted cells were suspended in debris removal solution (Miltenyi Biotec, #130-109-398) accordingly to manufacturer’s instructions. In some experiment, red erythrocytes were removed by incubation with red blood cell lysis buffer (abcam, #ab204733) for 3 minutes at room temperature. After washing, the cells were counted and resuspended in FBS containing 10% of DMSO (Applichem Panreac, #A3672.0050). The cell suspensions were aliquoted into freezing tubes (Nunc CryoTube, Thermo Fisher Scientific #366656) and placed in a CoolCell Cell Freezing container (Corning) at - 80°C. After at least 24 hours, the vials were transferred to liquid nitrogen for long-term storage.

### Staining for Flow cytometry

Fresh frozen vials of cell suspensions from patient samples listed (Extended Data Fig. 1a) were thawed by centrifugation at 350g for 7 min at 21°C in a 15mL falcon tube containing warm complete medium with 4 U/mL of benzonase (Merck, #70746-3). Complete medium consisted of RPMI-1640 medium (Thermo Fisher Scientific, #31870-025) supplemented with 5% of FBS (Thermo Fisher Scientific, #A5209402), 100 U/mL of Penicillin-Streptomycin (Thermo Fisher Scientific, #15140122), 2mM of L-Glutamine (Thermo Fisher Scientific, #25030024). Cell pellets were resuspended in complete medium containing 2 U/mL of benzonase and counted to ensure no more than 1.5 million cells were stained per sample to minimize variability between samples. The cells were transferred to a 96-well V-bottom plate, and staining was performed after washing the medium with PBS. Cell pellets were resuspended in 50 uL per well of PBS containing Zombie NIR Fixable Dye (Biolegend, #423106, dilution 1/600), Fc Receptor Blocking Solution (Biolegend, #422302, dilution 1/200), and True-Stain Monocyte Blocker (Biolegend, #426102, dilution 1/100), followed by incubation for 20 minutes at 4°C. After washing with PBS and centrifugation at 450g for 5 minutes, the supernatant was removed. Next, 50uL of RPMI-1640 medium supplemented with 3% of FBS at room temperature containing the antibody targeting chemokine receptor (CXCR6, CCR7, CX3CR1) was added, and the mixture was incubated for 15 minutes at 37°C. Subsequently, 50 uL of cold RPMI-1640 medium supplemented with 3% of FBS containing all remaining antibodies targeting extracellular proteins were added on top of each well and incubated for additional 30 minutes on ice. After washing with PBS and centrifuging at 450g for 5 minutes, the supernatant was removed. Fixation/Permeabilization buffer (Thermo Fisher Scientific, #00-5523-00; 150uL per well) was added and incubated for 35 minutes on ice. After centrifugation at 600g for 7 minutes, the supernatant was discareded. Next, 50uL of Permeabilization buffer (Thermo Fisher Scientific, #00-5523-00) containing antibodies targeting intracellular proteins (CTLA-4, EOMES, T-bet, Ki67, FoxP3 and TOX) was added to each well and incubated overnight at 4°C. The following morning, 50uL of Permeabilization buffer containing anti-TCF-1, anti-Granzyme B and streptavidin antibodies was added to each well and incubated for 35 minutes at 4°C. After a final wash with Permeabilzation buffer and centrifugation at 600g for 5 minutes, the supernatant was removed. The cells were resuspended in 100uL of PBS, and the acquisition was performed on the Cytek Aurora 5-laser flow cytometer and data were collected as FCS files. All antibodies used are listed in Extended Data Table 1.

### Rhapsody experiment

Fresh frozen vials of cell suspensions from patient samples listed (Extended Data Fig. 7a) were thawed by centrifugation at 250 g for 10 minutes at 21°C in a 15 mL Falcon tube containing warm complete medium. The complete medium consisted of RPMI-1640 medium (Thermo Fisher Scientific, #31870-025) supplemented with 10% FBS (Thermo Fisher Scientific, #A5670402). Cell pellets were resuspended in complete medium, transferred to FACS tubes, and centrifuged again. Cell pellets were resuspended in 100 µL of labeling buffer containing Zombie NIR Fixable Dye (BioLegend, #423106, dilution 1:600) and Fc Receptor Blocking Solution (BioLegend, #422302, dilution 1:200) and incubated for 10 minutes on ice. The labeling buffer consisted of PBS supplemented with 3% FBS and 0.1 g/L herring sperm DNA (Thermo Fisher Scientific, #15634-017). Next, 80 µL of labeling buffer containing anti-CD45 BV785, anti-CD38 BV650, anti-CD14 FITC (final dilution 1:100), anti-CD19 SuperBright 436 (final dilution 1:50), and anti-KLRG1 APC (final dilution 1:200) was added along with 20 µL of Sample Tag (BD Biosciences, #633781). After 30 minutes of incubation on ice, cells were washed with washing buffer (PBS supplemented with 3% FBS), centrifuged, and resuspended in sorting buffer. Sorting buffer consisted of HBSS (Thermo Fisher Scientific, #14175095) supplemented with 10% FBS and 10 mM HEPES (Thermo Fisher Scientific, #15630080). Cells were sorted as CD45-positive, live cells, SSC-A low, CD14-negative, CD19-negative using a BD FACSymphony™ S6 Cell Sorter with a 100 µm nozzle, sample agitation set at 300, 4-way purity, and maintained at 4°C. The sorted cells were pooled according to their sample tag index. After sorting, cells were centrifuged and resuspended in 100 µL of labeling buffer, to which 100 µL of AbSeq antibody mix was added. This mix contained pre-titrated AbSeq antibodies from the Discovery Panel (BD Biosciences, #625970) along with 14 additional AbSeq antibodies at 1:200 dilution, as detailed in the Extended Data Table 2. After two washes with PBS and one with sample buffer (BD Biosciences, #650000062), cells were resuspended in sample buffer, counted, and the cell concentration was adjusted to load no more than 50,000 cells per cartridge. Single-cell capture was performed according to the manufacturer’s instructions (BD Biosciences, Doc ID: 210967 Rev. 3.0). cDNA synthesis and template switching were carried out following the manufacturer’s instructions (BD Biosciences, Doc ID: 23-24020(03)). Library preparation from the beads was performed by the Functional Genomics Center Zurich, following the manufacturer’s instructions (BD Biosciences, Doc ID: 23-24020(03)). Sequencing was performed on the Illumina NovaSeq 6000 platform, with the following read configurations: 40,000 reads for whole transcriptome mRNA analysis, 1,000 reads for sample tags, 5,000 reads for TCR, and 25,000 reads for AbSeq. All antibodies and reagents used are listed in the Extended Data Table 2.

### Ex vivo stimulation

Fresh frozen vials of cell suspensions from the patient samples listed (Extended Data Fig. 8a) were thawed by centrifugation at 250 g for 10 minutes at 21°C in a 15 mL Falcon tube containing warm complete medium. The complete medium consisted of RPMI-1640 medium (Thermo Fisher Scientific, #31870-025) supplemented with 10% FBS (Thermo Fisher Scientific, #A5670402). Cell pellets were resuspended in 100 µL of live-dead labeling buffer containing Zombie NIR Fixable Dye (BioLegend, #423106, dilution 1:600) and Fc Receptor Blocking Solution (BioLegend, #422302, dilution 1:200) and incubated for 10 minutes on ice. The labeling buffer consisted of PBS supplemented with 3% FBS. Next, 100 µL of labeling buffer containing anti-CD45 PerCP, anti-CD3 BUV805, anti-CD4 Spark Blue 550, anti-CD8 PE, anti-CD56 PE-Dazzle594, anti-CD25 R718, anti-CD45RO PE-Cy7, CD161 BV785, CD127 BV605, CD103 BV510, and CD69 BV421 (information on clones and dilutions in Extended Data Table 3) were added to the well and incubated for 30 minutes on ice. After washing, cells were sorted as indicated: CD4 Mem1 (CD45RO^+^ CD127^+^ CD69^-^), CD4 Mem2 (CD45RO^+^ CD127^+^ CD69^+^ CD161^-^), CD8 Mem (CD45RO^+^ CD127^+^ CD69^-^) and CD8 EM (CD45RO^+^ CD127^-^ CD69^+^) according to the sorting strategy Extended Data Fig. 8b, using a BD FACSymphony™ S6 Cell Sorter with a 100 µm nozzle, sample agitation set to 300, 4-way purity, and maintained at 4°C.

Beforehand, 96-well plates (Sarstedt, #7510036) were prepared by coating with 0.5 µg/mL anti-CD3 antibody (clone OKT3, ThermoFisher Scientific, #16-0037-85) alone or combined with 1 µg/mL of Recombinant Human CD40-Fc Chimera recombinant protein (Biolegend, #777202) or anti-CD28 antibody (clone CD28.2, ThermoFisher Scientific, #16-0289-85). Each well was coated with 100 µL of PBS containing the antibody and incubated either for 2 hours at 37°C or overnight at 4°C, followed by three washes. After sorting, cells were centrifuged and resuspended in DMEM supplemented with 10% FBS, sodium pyruvate (1mM, ThermoFisher Scientific, #11360070), MEM Non-Essential Amina Acids (1X, ThermoFisher Scientific, #11140050), L-glutamine (2mM, ThermoFisher Scientific, #25030081), β-mercaptoethanol (AppliChem GmbH, # A1108,0100), and penicillin/streptomycin (Gibco, #7001592) to a concentration of 1 million cells/mL, based on the sorter count. After resting for 2 hours at 37°C, cells were counted, and the concentration was adjusted to load 30,000 cells in 100uL per well into the prepared 96-well plate Ab-coated. After 48 hours of incubation, cells were centrifuged, and the supernatant was stored at -80°C for LEGENDplex analysis. The cells were transferred to a V-bottom 96-well plate for activation marker staining.

Cells were stained with 50 µL of live-dead labeling buffer for 10 minutes on ice. Then, 50 µL of an antibody mix (anti-CD45 PerCP, anti-CD3 BUV805, anti-CD25 PE, CD69 BV421, CD27 BUV563, PD-1 BV605, CD28 BV750, TIM3 FITC, CD38 APC-Fire 810, HLA-DR BUV395) was added to the cell suspension and incubated for an additional 30 minutes at 4°C. Fixation/Permeabilization buffer (Thermo Fisher Scientific, #00-5523-00; 100 µL per well) was added and incubated for 35 minutes on ice. After washing, 100 µL of Permeabilization buffer (Thermo Fisher Scientific, #00-5523-00) containing antibodies targeting intracellular proteins (T-bet BV711, CD40L AF647, Granzyme B AF700, Granzyme K PE-Cy7, Ki67 BV480) was added to each well and incubated overnight at 4°C. The following morning, cells were washed with Permeabilization buffer and acquired on the Cytek Aurora 5-laser flow cytometer. All antibodies used are listed in the Extended Data Table 3. Cytokine measurement from the undiluted supernatant was performed using the LEGENDplex™ HU Th Cytokine Panel (12-plex, #741028, BioLegend) following the manufacturer’s instructions. Analysis was performed using the LEGENDplex data analysis software.

### Multiplex imaging on tissue microarrays

FFPE samples from the cohort in Extended Data Fig. 1a were H&E-stained, scanned, and a board-certified pathologist (B.S.) selected 600 µm tissue cores, five cores from each the tumor center and invasive margin compartment (as defined ^69^), and three from perilesional normal tissue, for construction of tissue microarrays (TMAs) using the TMA Grand Master (3D Histech). Sections of 2 µm were mounted on glass slides for re-evaluation and confirmation of region annotations. Of 390 cores, 17 were reclassified from tumor center to invasive margin, 13 from invasive margin to tumor center, and 80 were excluded due to disruption or uncertain regional assignment.

Sequential immunofluorescence imaging was performed using the COMET^TM^ platform (Lunaphore Technologies) using a 18-protein panel + DAPI. All antibodies used are listed in the Extended Data Table 4. Slides with FFPE tissue were pre-processed using the PreTreatment Module (PTM) and Deparaffinization and heat-induced epitope retrieval solution from Epredia^TM^ for the dewaxing and antigen-retrieval protocol (pH=9, Temperature= 102°C) performed on slides before the staining on COMET^TM^ as recommended by Lunaphore Technologies. The slides were then loaded onto the COMET^TM^ Stainer and processed according to manufacturer’s recommendations. Samples underwent iterative staining with primary antibodies, secondary antibodies, and subsequent imaging, followed by elution of the primary and secondary antibodies, and imaging of residual background, as previously described ^40^. All primary antibody incubations were performed for 4 to 8 minutes. The platform automatically processes the individual fields of view from each cycle, overlays, align and stitches them, resulting in a large multi-layer ome.tiff images file. Lunaphore Viewer was used to view the images and remove background fluorescence. QuPath 0.6.0-rc3 was used to define the TMA grid with the TMA dearrayer, identify and annotate disrupted regions, and perform cell segmentation using Instanseg ^70^, an embedding-based instance segmentation algorithm that uses DAPI and specified markers signals (CD3, CD20, CD11c, CD68 and CD56). After segmentation, each cell’s x and y coordinates and the median fluorescence signals from the nucleus and cell were exported for downstream R analysis.

### Analysis Flow Cytometry

After correction of the compensation matrix using FlowJo software (version 10.9.0), cells were gated on single, live immune cells (gating strategy shown in Extended Data Fig. 6a, up to “Cleaned CD45+”). Cleaned cells were exported as new FCS files and imported into R version 4.4.1 for analysis. Before automated high-dimensional data analysis, flow cytometry data were transformed using an inverse hyperbolic sine (arcsinh) function with a manually determined cofactor and normalized between 0 and 1 (Extended Data Table 5). A preliminary round of unsupervised clustering using FlowSOM (version 2.12.0) was conducted to clean the data and exclude non-lymphocyte cells from downstream analysis. Signal alignment for each marker across samples was checked. Samples with fewer than 1,000 cells were excluded from the analysis. For clustering of major lymphoid cells, UMAP was generated using 1,000 cells per sample and all markers except TIGIT, CD45, Live/Dead, and CD14. FlowSOM clustering was performed using CD3, CD4, CD8, CD161, CD45RO, CD127, CXCR6, CD56, KLRG1, TCRγδ, CD16, FoxP3, and CX3CR1 to identify 11 clusters. These clusters were subsequently merged based on marker expression into the following categories: Treg, CD4 T cells, CD8 T cells, MAIT, DP T cells, DN T cells, NK cells, TCRγδ T cells, and B cells.

For subsequent clustering of CD4 T cells, the previously identified major lymphoid CD4 T cell and Treg clusters were combined for downstream analysis. UMAP was generated using 1,000 cells per sample when possible. For UMAP and FlowSOM clustering, 24 markers were used, corresponding to all markers except TIGIT, CD45, Live/Dead, CD14, CD3, CD4, CD19, CD16, and TCRγδ. A total of 21 clusters were identified and merged based on marker expression into the following subsets: Naive, CM, Mem1, Mem2, RM, Mem CD161, SLEC, Effector, Effector PD1high, Tpex, Tex Term, Tex Prolif, T Prolif, and Treg. For subsequent clustering of CD8 T cells, UMAP was generated using 1,000 cells per sample when possible. For UMAP and FlowSOM clustering, 23 markers were used, corresponding to all markers except TIGIT, CD45, Live/Dead, CD14, CD3, CD4, CD8, CD19, CD16, and TCRγδ. A total of 10 clusters were identified and merged based on marker expression into the following subsets: Tex Term, RM, Tex Transitory, Innate-like, TEMRAs, SLEC, Effector, Mem, and Prolif. Subsequent analyses of all identified clusters were performed by merging the detailed CD4 and CD8 T cell clustering with other major lymphocyte populations, including NK cells, TCRγδ T cells, MAIT cells, DP T cells, DN T cells, and B cells.

Radar plots were generated using the *fmsb package* (version 0.7.6): For the phenotype of all 29 immune subsets identified in the merged analysis (Extended Data Figs. 2b and 3b), the axis limits for the radar plots are based on the minimum and maximum median values for each of the 29 lymphocyte markers across all 29 immune subsets. For the ex vivo stimulation (Fig. 4b,e), the radar plots were generated using the median fluorescence intensity. The axis limits for the radar plots are based on the minimum and maximum median values for each marker, determined across all groups (CD4 Mem1, CD4 Mem2, CD8 Mem, and CD8 EM) and under all stimulation conditions (unstimulated, aCD3/CD40, and aCD3/aCD28). For the 10 closest neighborhood spatial analysis (Fig. 5c and Extended Data Fig. 11b), radar plots show the mean cell type composition of CD4 or CD8 neighborhoods in the invasive margin and tumor center per response group. The radial axis of the radar plot is scaled to a maximum of 35%.

PCA analysis was performed using the *stats package* (version 4.4.1) to evaluate landscape changes by region based on the frequency of all 29 clusters across the 76 samples collected from all patients and grouped by region: 27 samples from the tumor center, 25 from the invasive margin, and 24 from the perilesional normal tissue (Extended Data Fig. 4c-d). To assess phenotypic changes by region, PCA analysis was performed on the median fluorescence intensity (MFI) of the 29 lymphocyte markers. As PCA analysis cannot handle NA values, clusters CD4 Tex Term, CD4 Tpex, and CD4 Tex Prolif, which were absent from several patients, were excluded. Additionally, the tumor center sample from patients 1 and 29, the invasive margin sample from patient 12, and the perilesional normal sample from patient 25, which were missing additional clusters, were removed. This analysis was performed on 26 immune subsets and a total of 25 samples from the tumor center, 24 from the invasive margin, and 23 from the perilesional normal tissue (Extended Data Fig. 4e,f). Finally, to evaluate changes in the immune landscape in response to NAC, PCA analysis was conducted by region based on immune subset frequency and grouped by TRG. After filtering on patient from which the TRG information was available, we depicted 22 patients in the tumor center (10 with TRG 4&5, 6 with TRG 3, and 6 with TRG 1&2), 21 patients in the invasive margin (10 with TRG 4&5, 6 with TRG 3, and 5 with TRG 1&2), and 20 patients in the perilesional normal tissue (11 with TRG 4&5, 4 with TRG 3, and 5 with TRG 1&2), grouped by TRG (Fig. 2e,f and Extended Data Fig. 5c-d).

### Module analysis

To interrogate the immune microenvironment, we examined co-enrichment patterns across the 29 identified lymphoid clusters from the 76 samples. Spearman correlations between lymphoid subset frequencies across all samples were calculated, followed by hierarchical clustering, which identified three distinct cell modules with coordinated behavior.

To calculate the abundance of each module, we performed the following steps:

1. Equal weighting for each cell type was ensured by normalizing the frequency of each cluster per sample between 0 and 1 to the 1st–99th percentile.
2. Module scores were calculated as the mean frequency of all immune subsets within each module (i.e., sum of frequencies divided by the number of cell types per module). This ensured that scores ranged from 0 to 1, giving equal importance to each module regardless of size.
3. The module with the highest score was identified as the most abundant module per sample.
4. The scores were transformed into percentages representing the abundance of each module per patient. This normalization ensured that the sum of module abundances for each patient equaled 100%.
5. Module abundance was evaluated across all samples by region.

Notably, module analysis was also performed within the tumor center, and metastatic module abundance was evaluated by TRG. For this analysis, tumor center samples were filtered, and the module analysis was repeated following the same steps as above. However, the frequencies of each lymphocyte subset were not normalized again, preserving the coherence of the general composition per region.

### Rhapsody analysis

#### Data processing, QC and cleaning

The BD Rhapsody Sequence Analysis v1.12.1 pipeline was used to align reads to the human reference genome (Build GRCh38.p13 with gene model definition from GENCODE, release 37) and generate the gene-cell unique molecular identifier (UMI)-adjusted matrix (RSEC) for each sample. The obtained raw count matrix was loaded into the Seurat workflow (version 5.1.0) for downstream analyses. Genes detected in fewer than three cells in each cartridge were filtered out prior to analysis. Quality control (QC) of cells was performed using three sequential metrics: total counts, the number of detected genes, and the proportion of mitochondrial gene counts per cell. Cells with mitochondrial gene expression exceeding 24% were excluded, as well as cells with fewer than 800 detected genes. To eliminate potential doublets, cells with total counts over 15,000 and detected genes exceeding 4,000 were also removed. For multiplexing, following expert recommendations from the BD Rhapsody platform, multiplets were recalled when a tag represented more than 75% of the total tag reads. After QC, the data matrix was normalized to 10,000 counts per cell using *Seurat::NormalizeData()* function, ensuring comparability among cells. The normalized counts were log-transformed as part of this process, and data scaling was performed using *Seurat::ScaleData()* function. Highly variable genes were identified using the *Seurat::FindVariableFeatures()* function with the variance-stabilizing transformation (VST) method, selecting the top 2,000 most variable genes for downstream analysis. Principal Component Analysis (PCA) was performed on these highly variable genes using the *Seurat::RunPCA()* function. A shared nearest neighbor (SNN) graph was constructed using *Seurat::FindNeighbors()* function, based on the first 13 PCs from the PCA space. Cells were clustered into distinct groups using *Seurat::FindClusters()* function, employing the Louvain algorithm. Based on mRNA clustering, four clusters that were not pure NK or T cells were removed from subsequent analyses. For Antibody-Derived Tag (ADT) data, variable features were defined using all features within the ADT assay. Normalization was performed using the Centered Log Ratio (CLR) transformation with *Seurat::NormalizeData()* function, normalizing across features (margin = 2). The normalized data were scaled using *Seurat::ScaleData()*, and PCA was performed with *Seurat::RunPCA()* function. Using the first 10 PCs, an SNN graph was constructed with *Seurat::FindNeighbors()* function, and clustering was conducted using *Seurat::FindClusters().* Clusters with inconsistent staining, reflecting potential doublets, and clusters that did not represent NK or T cells were removed from the analysis.

#### Identification of CD4 and CD8 T cell

For downstream analysis, both the RNA and ADT assays were normalized and scaled again and the cleaned data, as described before. As each patient were received in different days and the three regions were processed independently, the RNA data set was corrected for batch effects and integrate across patients using harmony integration ^34^. This was performed using *Seurat::RunHarmony()* function (lambda = 1.5 and theta = 3). Fourteen harmony clusters were identified based on RNA clustering analysis using the first 12 PC and a resolution of 0.9. Similarly, the ADT data set was corrected for batch effects and integrated using Seurat’s anchoring method ^71^. Features such as CD14, CD19, IgD, and IgM were excluded due to their lack of relevance. Integration anchors were identified using the *Seurat::FindIntegrationAnchors()* function. The datasets were integrated using the *Seurat::IntegrateData()* function, and the resulting integrated assay was stored as IADT in the original dataset. Five major populations were identified based on IADT clustering analysis, utilizing CD3, CD8, CD56, CD4, CD161, KLRG1, CD127, CD45RO, and TCRγδ as protein markers for clustering, with a resolution of 0.2. The identified populations were CD4 T cells, CD8 T cells, NK cells, TCRγδ T cells, and MAIT cells. To study CD4 and CD8 T cells separately, we implemented a multistep filtering process. First, CD4 and CD8 T cells were identified based on clusters in the integrated proteomic dataset (IADT). To refine these clusters, we reassigned CD4 Treg cells identified from the integrated RNA clustering analysis to the CD4 T cell population, excluding any CD8 T cells and ensuring that all Treg were identified. Finally, clusters corresponding to MAIT cells, NK cells, and stressed cells, as identified from the integrated RNA clustering analysis, were excluded from both the CD4 and CD8 T cell populations.

#### Analysis CD4 and CD8 T cell

Both CD4 and CD8 T cell RNA data set were scaled again and the proteomic IADT and transcriptomic harmony integrated modalities were integrated using the Weighted Nearest Neighbor (WNN) workflow ^72^ using *Seurat::FindMultiModalNeighbors()* function with PC 13 of the harmony RNA and PC 10 of the IADT. For CD4 T cells, resolution 0.5 was used to identify 10 clusters, two clusters were merged as Mem1 and two as Treg; while for CD8 T cells, resolution 0.4 was used to identify 8 clusters. For visualization, the dimensionality was further reduced using UMAP implemented in the *Seurat::RunUMAP()* function. Differentially expressed genes were identified using the *Seurat::FindAllMarkers()* function, reporting only genes with positive log fold change, a minimum detection rate of 25%, and a log fold change threshold of 0.25 (Extended Data Table 6 and 7). Rhapsody signature of each subset was defined by filtering differentially expressed genes that are detected in more than 30% of cells in the target cluster (pct.1 > 0.3), less than 30% of cells in other clusters (pct.2 < 0.3), and if their average log2 fold change (avg_log2FC) exceeded 0.3 (Extended Data Table 8).

Signatures z-score were calculated using the function *Seurat::AddModuleScore()* using list of genes from previously described signatures: stem-like memory signature ^55^ for CD8 T cells and ^66^ for CD4 T cells, circulating memory and residency T cell signatures ^73^, cytotoxic CD8 ^74^ T cells signatures, and MSigDB signatures for TGFb, Hypoxia. Th1 and Th17 signatures were defined using curated gene sets, with Th17 characterized by AHR, CCR6, TGFB1, REL, RELA, IL23R, CCL20, CEBPD, STAT3, STAT4, and IL23A; and Th1 defined by IFNG, XCL1, TBX21, CCL4, CCL5, CCL3, CSF2, IL2, STAT1, CXCR3, and TNF.

Single-cell regulatory network inference and clustering (SCENIC) on CD4 T cells was performed using the pySCENIC workflow ^31^. Genes expressed in less than 0.5% of cells were filtered out, and the rest was performed with standard parameters. The reference files used were allTFs_hg38, hg19-tss-centered-10kb-7species.mc9nr.genes_vs_motifs.rankings.feather and motifs-v9-nr.hgnc-m0.001-o0.0.tbl, downloaded from the aertslab resources cistarget databases and motif2tf.

TCR sequencing data were analyzed using the scRepertoire (v2.2.1) workflow ^75^ and merging all cluster from the CD4 and CD8 T cell analysis with WNN workflow clustering. We used the *clonalBias()* function ^76^ to assess clonal skewing across clusters by patient and calculating baseline frequencies with a minimum expansion threshold of five. We used the *clonalOverlap()* function with the "morisita" method to evaluate and visualize the similarity of TCR clonotypes across clusters.

### Data mining Liu et al., 2022

The data set from Liu et al., 2022 ^23^ was downloaded and included 10 patients with CRLM, who had not undergone NAC treatment. We used the major annotation from the original dataset to extract the CD4 T cell cluster, comprising 13,416 cells from the tumor center tissue (metastatic tumor) and 4,551 cells from the perilesional normal tissue (metastatic normal). Harmony was used to integrate this dataset with the CD4 T cells from our dataset, which included 15,470 cells, to correct for both batch effects and patient effects, as the samples were processed independently. This integration was performed using *Seurat::RunHarmony()* function (lambda = 1.5, theta = c(3,3)) on the variable features from the Rhapsody data. Seven clusters were identified after integration, based on RNA clustering analysis using the first 12 PCs and a resolution of 0.4. Similarly, we used the major annotation from the original dataset to extract the CD8 T cell cluster, which included 7,198 cells from the tumor center tissue (metastatic tumor) and 15,248 cells from the perilesional normal tissue (metastatic normal). Harmony was used again the same parameters than before, to integrate this dataset with the CD8 T cells from our dataset, containing 19,767 cells, to correct for both batch effects and patient effects, as the samples were processed independently. Nine integrated clusters were identified based on RNA clustering analysis using the first 12 PCs and a resolution of 0.5.

For visualization, dimensionality was further reduced using UMAP, implemented with the *Seurat::RunUMAP()* function. Signature z-scores were computed in the untreated patient dataset using the *Seurat::AddModuleScore()* function, with gene lists from the previously described Rhapsody signature.

### Data mining Wu et al., 2022

The data set from Wu et al., 2022 ^35^ was downloaded and included 20 patients with CRC Stage IV: 11 with CRLM and no treatment (pretreated, 21,423 cells), six patients treated with NAC with progressive disease or stable disease (treated-PD/SD, 7,167 cells), and three patients treated with NAC with partial response (treated-PR, 20,671 cells). The raw count matrix was loaded into the Seurat workflow for downstream analyses. Quality control (QC) of cells was performed using three sequential metrics: total counts, the number of detected genes, and the proportion of mitochondrial gene counts per cell. Cells with mitochondrial gene expression exceeding 10% were excluded, as were cells with fewer than 400 detected genes. To eliminate potential doublets, cells with total counts over 15,000 and detected genes exceeding 3,000 were also removed. Batch effect per patient sample was correct using *Seurat::RunHarmony()* function (lambda = 1.5, theta = 3).

We identified sixteen clusters based on RNA clustering analysis using the first 12 PCs and a resolution of 0.9. Six of these clusters were identified as CD4 T cells, including Tregs, and were reclustered following the previously described method. After re-clustering on CD4 T cells using the first 10 PCs and a resolution of 0.4, we identified six clusters. For visualization, dimensionality was further reduced using UMAP, implemented with the *Seurat::RunUMAP()* function. Signature z-scores were calculated using the *Seurat::AddModuleScore()* function with the following gene lists:

- Circulating, less differentiated memory: S1PR1, CCR7, KLF3, CD55
- Tissue Th1-like: CD69, IFNG, CD40LG, TNF, TBX21
- Th17-like: KLRB1, AHR, CCL20, IL23R
- Suppressive Treg: FOXP3, IL2RA, CCR8, IL1R1

### Analysis of Multiplex-imaging

Before automated high-dimensional data analysis, multiplex-imaging data were transformed using an inverse hyperbolic sine (arcsinh) function with a manually determined cofactor (Extended Data Table 9) and normalized between 0 and 1. Batch correction between the four slides of TMAs was performed using cyCombine algorithm ^77^ with batch as slide and considering the region (tumor center, invasive margin and perilesional normal tissue) as covariable. A preliminary round of unsupervised clustering using FlowSOM (version 2.12.0) was conducted to clean the data from artefact and exclude unstained cells (CD3, HLA-DR, CD4, CD56, CD8, CD11c, CD68, CD20, SATB2, PanCK, CD69, CD11b) from downstream analysis. The resulting cells were first annotated as immune cells (expressing CD3, CD4, CD56, CD8, CD20, HLA-DR, CD11c, CD68 or CD11b), non-tumor epithelial cells (expressing PanCK but SATB2 negative), cholangiocytes (expressing PanCK and CD56) or tumor cells (expressing both PanCK and SATB2) using 15 clusters from FlowSOM clustering based on the previously mentioned markers. For subsequent clustering, the data for the immune cells were transformed and batch corrected again as previously described and in order to improve the signal separation on this compartment and cells with the DAPI nucleus signal lower than 250 or an area lower than 15µm^2^ were filtered out. Immune cells were then subclustered using CD3, CD4, CD56, CD8 and CD20 for FlowSOM to identify 7 clusters which were merged into the following categories: T cells, B cells, Myeloid cells and NK cells. T cells were then subclustered with FlowsSOM to ientify 4 clusters namely CD4 T cells, CD8 T cells, other T cells (CD3^+^ CD4^-^ CD8^-^) and Tregs. The identification of Treg cells was optimized by considering CD4 T cells with FOXP3 signal higher than 0.25, CD4 signal higher than 0.15 and CD8 signal lower than 0.35. Finally, CD4 T cells were subclustered using TCF1, Ki67, CD69 and PD-1 to identify 15 clusters merged into CD4 Tex Term (TCF1^-^ PD1^+^), CD4 Tpex (TCF1^+^ PD-1^+^), CD4 prolif (Ki67^+^), CD4 TCF1^+^ CD69^-^, CD4 TCF1^+^ CD69^+^, CD4 TCF1^-^ CD69^+^ and CD4 TCF1^-^ CD69^-^. CD8 T cells were subclustered using TCF1, Ki67, GzmK and PD-1 to identify 18 clusters merged into CD8 Tex Term (TCF1^-^ PD1^+^), CD8 prolif (Ki67^+^), CD8 TCF1^+^ GzmK^-^, CD8 TCF1^+^ GzmK ^+^, CD8 TCF1^-^ GzmK ^+^ and CD8 TCF1^-^ GzmK ^-^. Myeloid cells were subclusterd using CD68, HLA-DR, CD11c, CD11b to identify 9 clusters merged into DCs (CD68^-^ HLA-DR^+^ CD11c^+^), Macrophages (CD68^+^) HLA-DR^+^ CD11c^+^ or HLA-DR^+^ CD11c^-^ and myeloid CD11b^+^ HLA-DR^+^ or CD11b^+^ HLA-DR^-^. UMAP was generated using all the annotated cells (408’311) based on CD3, HLA-DR, CD4, CD56, CD8, CD11c, CD68, CD20, SATB2, PanCK, FOXP3, CD11b. Density of the different subsets was calculated by dividing the sum of cell count of all TMAs collected by the sum of the area (mm^2^), for each patient and region. The area was calculated using the tissue detection feature in QuPath (Threshold = 12; requested pixel size = 2; max fill area = 10000; dark background mode; smoothing image / coordinate and cleanup with median filter enabled) from which we subtracted areas being annotated as disrupted.

To investigate spatial interactions between immune cell populations, we calculated pairwise Euclidean distances between all cells within each individual TMA core. For downstream quantification, we filtered for distance occurring within a 15 μm threshold, representing putative physical proximity for cell–cell communication. For each TMA punch, we computed the number of close-range cell–cell proximities for all cell type pairs, we generated all possible combinations of cell type pairs across all TMA punches and filled in zero values for missing combinations. Interaction counts were normalized to the segmented tissue area (yielding a density of interactions per mm²) for each patient and region. The median interaction density for responder and non-responder is displayed.

Neighborhood analysis of CD4 T cells in CRLM tissues was performed by retrieving the 10 closest non-CD4 T cell neighbors to each CD4 T cell. For each patient and region, we computed the neighbor cell type composition and summarized each response group by the mean proportion. For visualization, we showed the profiles in a radar plot (fmsb::radarchart) for responders (TRG1&2) and non-responders (TRG4&5), with a radial axis to a maximum of 35%. Neighborhood analysis of CD8 T cells in CRLM tissues was performed in the same manner, retaining the 10 closest non-CD8 T cell neighbors to each CD8 T cell.

The proximity score was calculated as the ratio of identified interactions (using a 15 µm radius threshold to CD4 or CD8 T cells) to the total number of cells of the interacting type, providing insight into how frequently these cell type interacts relative to their population size ^78^. Scores were averaged across blocks within the same responder group.

To identify spatially proximate immune cell triads We built an undirected proximity network per sample and region from all cell pairs with Euclidean distance 0<d<15μm. We identified fully connected subgraphs (cliques) of size ≥3 and retained only those containing at least one CD4 T cell, one stem-like CD8 T cell (TCF1⁺ GzmK⁺ or TCF1⁺ GzmK⁻), and one antigen-presenting cell (B cells, dendritic cells, macrophages HLA-DR⁺ CD11c⁺, macrophages HLA-DR⁺, or myeloid CD11b⁺ HLA-DR⁺). We then counted the number of cells participating in at least one qualifying triad and normalized by the segmented tissue area to report cells-in-triads per mm². Finally, for APC-type in triad, we computed the density of APC triad members per subset and summarized these per-subset densities by response group using the mean across patients.

### Statistical analysis

In the Flow Cytometry analysis, the Rhapsody, the data mining analyses and the multiplex-analysis, statistical significance in frequency or density was determined using paired (when comparing regions) or unpaired (when comparing response group) Wilcoxon tests corrected for multiple comparisons using the Benjamini-Hochberg method. The analysis was conducted in R with the *compare_means()* function from the ggpubr package (version 0.6.0). Significance levels are displayed as follows, * pvalue adjusted <0.05, ** <0.01, *** < 0.001,**** <0.0001. We calculated the effect size of changes of MFI using Cohen’s d, the calculation was performed in R using the effsize package (version 0.8.1), specifically the *cohen.d()* function Spearman correlation test was used to measure the strength and direction of the frequency of each immune subset to the TRG. A positive Spearman correlation coefficient of Frequency vs TRG indicates an association with low TRG (i.e. responders), while a negative correlation indicates association with high TRG (i.e. non-responder). Significance levels are displayed as follows, * pvalue adjusted <0.05, ** <0.01, *** < 0.001,**** <0.0001.

For the scenic analysis, Wilcoxon rank-sum test was used to identify differentially active regulons between memory clusters (CD4 Mem1, CD4 Mem2 and CD4 Mem CD161). For each regulon with sufficient representation (more than 50 cells in the target cluster), the test compares the regulon activity values in the target cluster against the other clusters, calculating a p-value and log fold change for the difference in activity. The p-values are adjusted for multiple testing using the Benjamini-Hochberg (BH) method.

For statistical analysis of the 10 closest neighbor proportions between response groups (responder vs. non-responder, mid vs. non-responder, and responder vs. mid) in the multiplex imaging data, we used non-parametric permutation-based independence tests (*independence_test()* function from the coin package (version 1.4.3)). P-values were corrected for multiple testing using the Benjamini–Hochberg method. Significance thresholds were indicated as follows: * pvalue adjusted <0.05, ** <0.01. Statistical testing for the proximity score, was performed separately for each neighbor type and region, an initial Kruskal–Wallis test was performed followed by Benjamini–Hochberg-adjusted pairwise Wilcoxon rank-sum tests for all group pairs (responder, mid, non-responder). Neighbor types were included only if each group contained at least two samples and neither group exhibited zero variance in scores.

Triad densities were compared by region across responder groups using Wilcoxon rank-sum tests stratified by anatomical region using the function (ggpubr::compare_means). Significance pvalue were reported as follows: * <0.05, ** <0.01, *** < 0.001. The APC-type in triad were compared using two sided Wilcoxon test with the function (stats::wilcox.test) applied within each combination of cluster, region, and responder group. Statistical significance was annotated as follows: * pvalue <0.05 and ** <0.01.

## Notes

### Competing Interest Statement

The authors have declared no competing interest.

### Summary of Updates

The text was improved, we also merged figure 1 and 2 and added 1 graph to figure S9. Finally, one grant was added in the finance as we initially forgot to add it.

