## Supplementary material for "CD4 memory T cells orchestrate therapy-responsive immune niches in colorectal cancer liver metastases": Document 1. Fig. S1- S11, Tables S1-S5, S9

### Extended Data

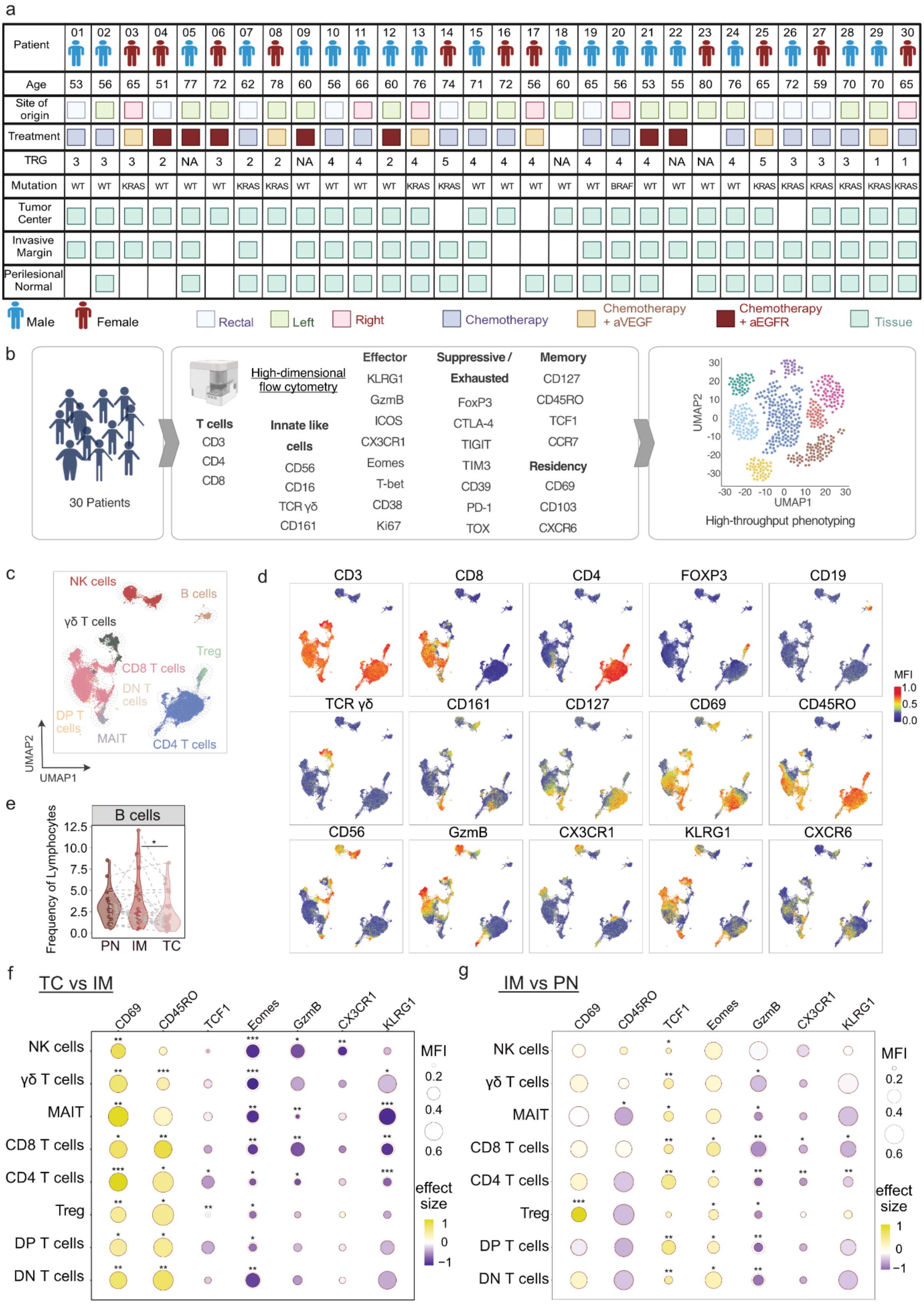
a
Patient
01 02 03 04 05 06 07 08 09 10 11 12 13 14 15 16 17 18 19 20 21 22 23 24 25 26 27 28 29 30
Age
53 56 65 51 77 72 62 78 60 56 66 60 76 74 71 72 56 60 65 56 53 55 80 76 65 72 59 70 70 65
Site of origin
Treatment
TRG
3 3 3 2 NA 3 2 2 NA 4 4 2 4 5 4 4 4 NA 4 4 4 NA NA 4 5 3 3 3 1 1
Mutation
WT WT KRAS WT WT WT KRAS KRAS WT WT WT WT KRAS KRAS WT WT WT WT WT BRAF WT WT WT WT KRAS KRAS KRAS KRAS KRAS KRAS
Tumor Center
Invasive Margin
Perilesional Normal
Male
Female
Rectal
Left
Right
Chemotherapy
Chemotherapy + aVEGF
Chemotherapy + aEGFR
Tissue
b
30 Patients
High-dimensional flow cytometry
T cells
CD3
CD4
CD8
Innate like cells
CD56
CD16
TCR γδ
CD161
Effector
KLRG1
GzmB
ICOS
CX3CR1
Eomes
T-bet
CD38
Ki67
Suppressive / Exhausted
FoxP3
CTLA-4
TIGIT
TIM3
CD39
PD-1
TOX
Memory
CD127
CD45RO
TCF1
CCR7
Residency
CD69
CD103
CXCR6
UMAP2
UMAP1
High-throughput phenotyping
c
NK cells
B cells
γδ T cells
Treg
CD8 T cells
DN T cells
DP T cells
MAIT
CD4 T cells
UMAP2
UMAP1
d
CD3
CD8
CD4
FOXP3
CD19
TCR γδ
CD161
CD127
CD69
CD45RO
CD56
GzmB
CX3CR1
KLRG1
CXCR6
MFI
1.0
0.5
0.0
e
B cells
Frequency of Lymphocytes
12.5
10.0
7.5
5.0
2.5
0.0
PN
IM
TC
f
TC vs IM
g
IM vs PN
CD69
CD45RO
TCF1
Eomes
GzmB
CX3CR1
KLRG1
NK cells
γδ T cells
MAIT
CD8 T cells
CD4 T cells
Treg
DP T cells
DN T cells
MFI
0.2
0.4
0.6
effect size
1
0
−1

**Extended Data Fig. 1: The tumor center in CRLM is enriched with CD4 memory T cells and exhibits reduced cytotoxic activity.** (a) Clinical information across 75 samples from 30 patients used for high-dimensional flow cytometry (NA, not assessable). (b) Schematic of the high-dimensional flow cytometry analysis. (c) UMAPs of all major lymphoid population colored by cluster. (d) UMAPs of all major lymphoid population colored by normalized marker intensity. (e) Violin plot showing B cell frequencies across regions. (f) Bubble plot comparing median normalized marker expression in tumor center (TC, darker circle) vs invasive margin (IM, lighter circle) and colored by effect size (Cohen's D), indicating upregulation (yellow) or downregulation (blue) of marker expression. (g) Bubble plot comparing median normalized marker expression in invasive margin (IM, darker circle) vs perilesional normal tissue (PN, lighter circle) and colored by effect size (Cohen's D), indicating upregulation (yellow) or downregulation (blue) of marker. (e, f, g) Statistical significance was determined by paired Wilcoxon tests with Benjamini-Hochberg correction. Adjusted P values are shown as *P <0.05, **P <0.01, ***P < 0.001,****P <0.0001.

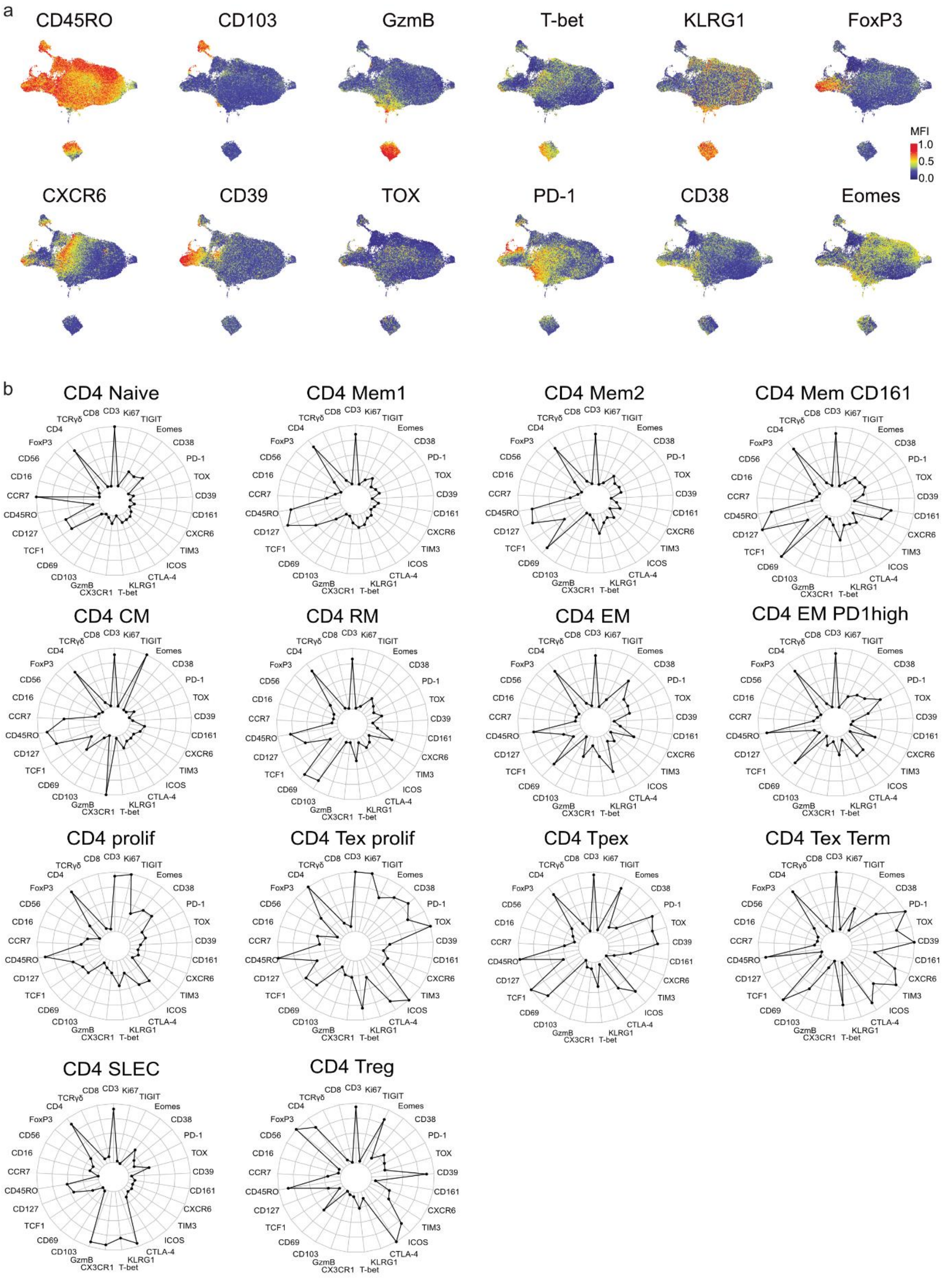
a
CD45RO
CD103
GzmB
T-bet
KLRG1
FoxP3
MFI
1.0
0.5
0.0
CXCR6
CD39
TOX
PD-1
CD38
Eomes
b
CD4 Naive
CD4 Mem1
CD4 Mem2
CD4 Mem CD161
CD4 CM
CD4 RM
CD4 EM
CD4 EM PD1high
CD4 prolif
CD4 Tex prolif
CD4 Tpex
CD4 Tex Term
CD4 SLEC
CD4 Treg
CD3
Ki67
TIGIT
Eomes
CD38
PD-1
TOX
CD39
CD161
CXCR6
TIM3
ICOS
CTLA-4
KLRG1
T-bet
CX3CR1
GzmB
CD103
CD69
TCF1
CD127
CD45RO
CCR7
CD16
CD56
FoxP3
CD4
TCRγδ
CD8

**Extended Data Fig. 2: Phenotypic characterization of the CD4 T cells in CRLM.** (a) UMAPs of CD4 T cells colored by normalized marker intensity. (b) Radar plots showing median fluorescence intensity profiles of 29 features across all lymphocytes for each CD4 T cell cluster.

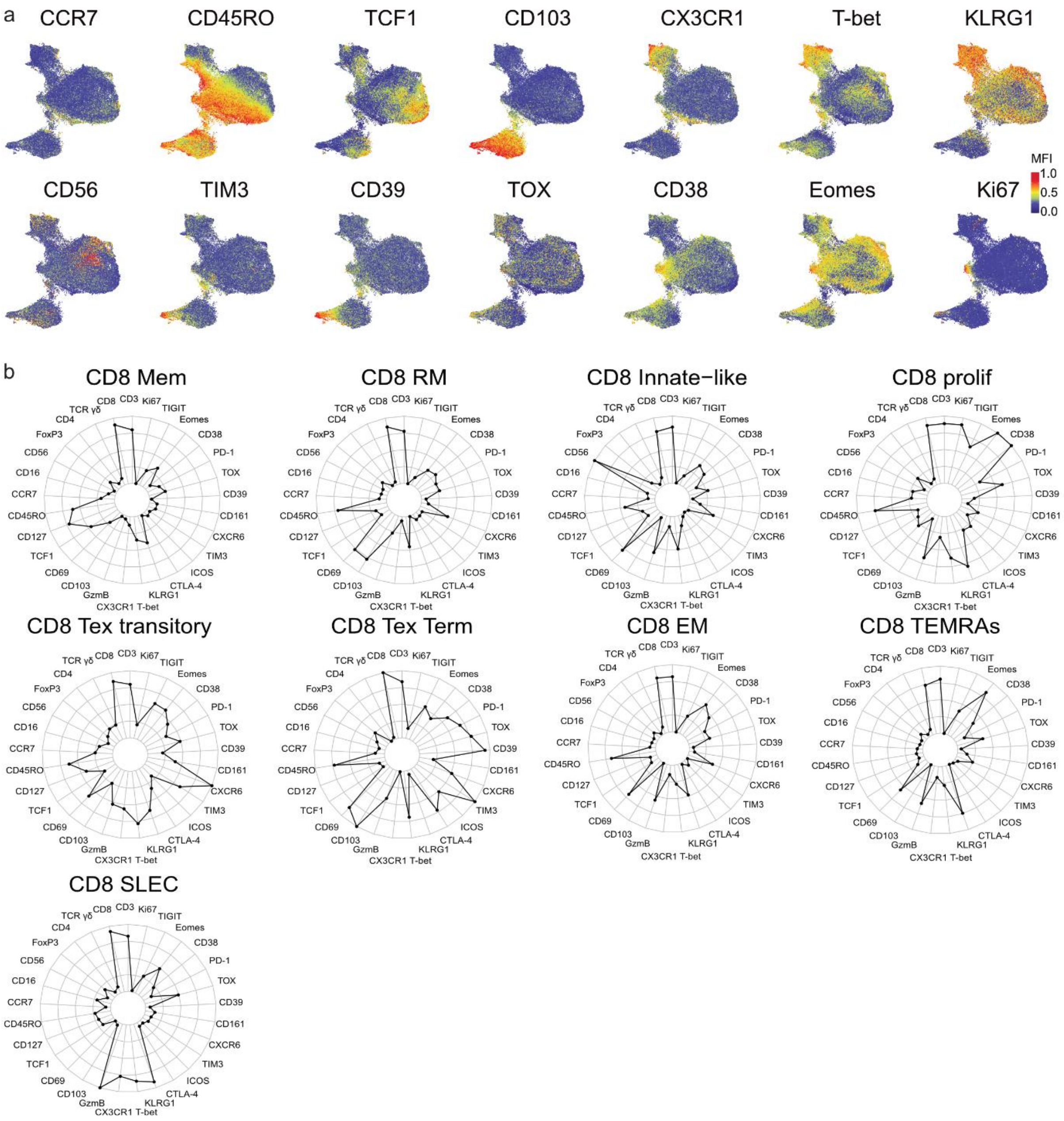

a
CCR7
CD45RO
TCF1
CD103
CX3CR1
T-bet
KLRG1
MFI
1.0
0.5
0.0
CD56
TIM3
CD39
TOX
CD38
Eomes
Ki67
b
CD8 Mem
CD8 RM
CD8 Innate−like
CD8 prolif
CD8 Tex transitory
CD8 Tex Term
CD8 EM
CD8 TEMRAs
CD8 SLEC
CD3
Ki67
TIGIT
Eomes
CD38
PD-1
TOX
CD39
CD161
CXCR6
TIM3
ICOS
CTLA-4
KLRG1
CX3CR1 T-bet
GzmB
CD103
CD69
TCF1
CD127
CD45RO
CCR7
CD16
CD56
FoxP3
CD4
TCR γδ
CD8

**Extended Data Fig. 3: Phenotypic characterization of the CD8 T cells in CRLM.** (a) UMAPs of CD8 T cells colored by normalized marker intensity. (b) Radar plots showing the median fluorescence intensity profiles of 29 features across all lymphocytes for each CD8 T cell cluster.

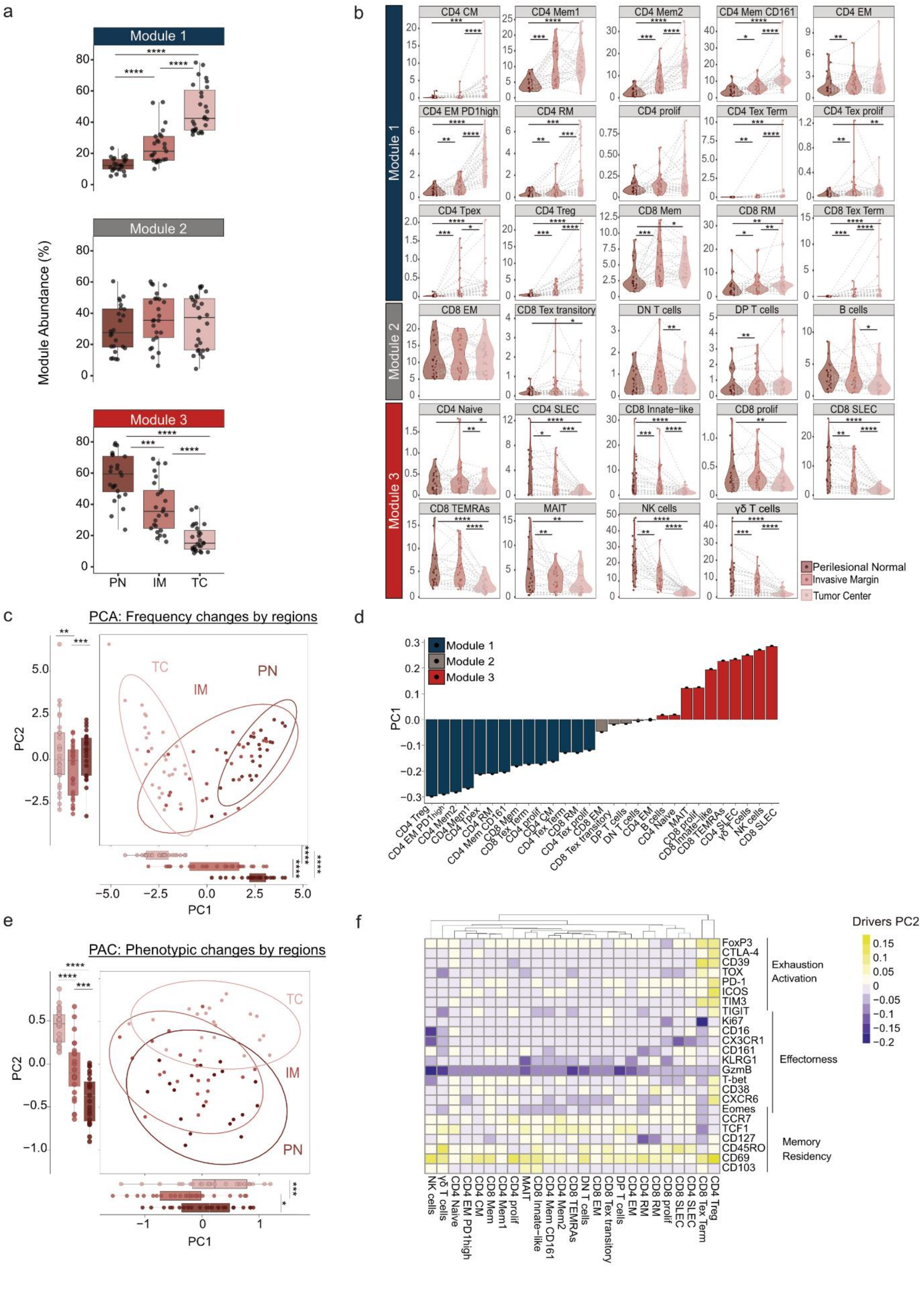

a
Module 1
Module 2
Module 3
Module Abundance (%)
PN
IM
TC
b
CD4 CM
CD4 Mem1
CD4 Mem2
CD4 Mem CD161
CD4 EM
CD4 EM PD1high
CD4 RM
CD4 prolif
CD4 Tex Term
CD4 Tex prolif
CD4 Tpex
CD4 Treg
CD8 Mem
CD8 RM
CD8 Tex Term
CD8 EM
CD8 Tex transitory
DN T cells
DP T cells
B cells
CD4 Naive
CD4 SLEC
CD8 Innate-like
CD8 prolif
CD8 SLEC
CD8 TEMRAs
MAIT
NK cells
γδ T cells
Perilesional Normal
Invasive Margin
Tumor Center
c
PCA: Frequency changes by regions
PC1
PC2
d
Module 1
Module 2
Module 3
PC1
e
PAC: Phenotypic changes by regions
f
Drivers PC2
FoxP3
CTLA-4
CD39
TOX
PD-1
ICOS
TIM3
TIGIT
Ki67
CD16
CX3CR1
CD161
KLRG1
GzmB
T-bet
CD38
CXCR6
Eomes
CCR7
TCF1
CD127
CD45RO
CD69
CD103
Exhaustion
Activation
Effectorness
Memory
Residency

**Extended Data Fig. 4: Spatial remodeling of the lymphoid landscape reveals enrichment of memory, exhausted, and suppressive T cells in the tumor center.** (a) Abundance of the three identified cellular modules across regions for all patients. (b) Violin plot showing frequencies of 29 lymphocytes subsets grouped by modules across perilesional normal (PN), invasive margin (IM), and tumor center (TC) regions. (c) PCA of immune subset frequencies, where each dot represents a patient sample from a given region, showing PC1 and PC2 separation across regions. Box plots display the median and interquartile range of PC1 and PC2 values per region. (d) PC1 distance of immune subsets separating TC from PN, colored by module. (e) PCA of phenotypic changes using median marker expression across all 76 samples from 30 patients, grouped by region. Boxplots show median and interquartile range of PC1 and PC2 values. (f) Heatmap of PC2 distances for each marker across subsets, depicting phenotypic changes by regions. (a, b, c, e) Statistical significance was assessed using paired Wilcoxon tests with Benjamini-Hochberg correction. Adjusted P values are indicated as *P <0.05, **P <0.01, ***P < 0.001,****P <0.0001.

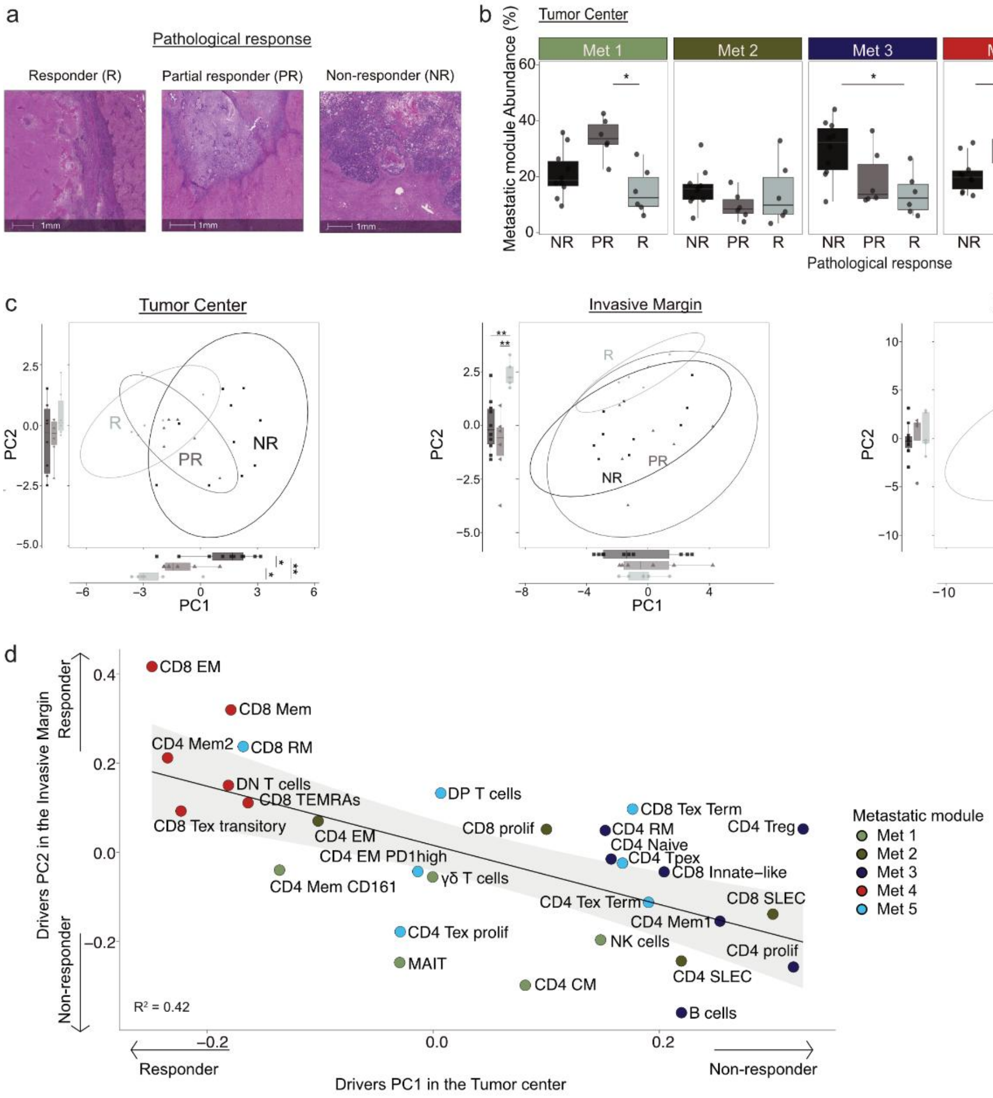
a
Pathological response
Responder (R)
Partial responder (PR)
Non-responder (NR)
1mm
b
Tumor Center
Metastatic module Abundance (%)
Met 1
Met 2
Met 3
Met 4
Met 5
NR PR R
Pathological response
c
Tumor Center
Invasive Margin
Perilesional Normal
PC2
PC1
R
PR
NR
d
CD8 EM
CD8 Mem
CD4 Mem2
CD8 RM
DN T cells
CD8 TEMRAs
CD8 Tex transitory
DP T cells
CD4 EM
CD8 prolif
CD8 Tex Term
CD4 Treg
CD4 RM
CD4 Naive
CD4 Tpex
CD4 EM PD1high
CD8 Innate−like
CD4 Mem CD161
γδ T cells
CD4 Tex Term
CD8 SLEC
CD4 Mem1
CD4 Tex prolif
NK cells
CD4 prolif
MAIT
CD4 SLEC
CD4 CM
B cells
R² = 0.42
Drivers PC2 in the Invasive Margin
Responder
Non-responder
Drivers PC1 in the Tumor center
Metastatic module
Met 1
Met 2
Met 3
Met 4
Met 5

**Extended Data Fig. 5: Response to neoadjuvant therapy is associated with a distinct module of CD4 and CD8 memory and effector cells in the tumor center and invasive margin.** (a) Representative H&E staining from responders (R, TRG 1&2), partial responders (PR, TRG 3), and non-responders (NR, TRG 4&5). (b) Abundance of metastatic module across patients stratified by pathological response. (c) PCA of immune subset frequencies, with each dot representing one patient sample per region (n=22 tumor center, n= 21 invasive margin, n= 20 perilesional normal). Patients are grouped by response: R (circles), PR (triangles), and NR (squares). Boxplots show median and interquartile range of PC1 and PC2 values for each group. (d) PC1 distance (tumor center) and PC2 distance (invasive margin) for each immune subset, colored by metastatic module (defined in Fig. 2a). Linear correlation is indicated ($R^2$ = 0.42). (b, c) Statistical significance was assessed with unpaired Wilcoxon tests and Benjamini-Hochberg correction. Adjusted P values are shown as *P <0.05, **P <0.01.

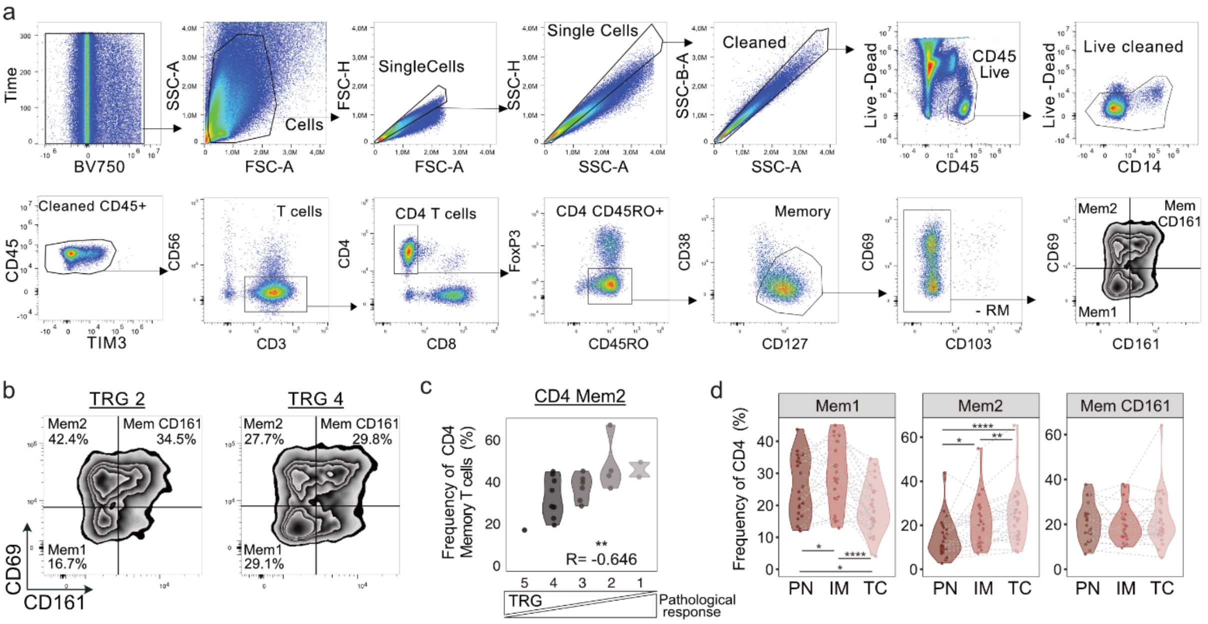

a
Time
BV750
SSC-A
FSC-A
Cells
FSC-H
SingleCells
SSC-H
Single Cells
SSC-B-A
Cleaned
Live -Dead
CD45 Live
CD45
Live cleaned
CD14
Cleaned CD45+
CD45
TIM3
CD56
T cells
CD3
CD4 T cells
CD4
CD8
FoxP3
CD4 CD45RO+
CD45RO
CD38
Memory
CD127
CD69
- RM
CD103
Mem2
Mem CD161
Mem1
CD161
b
TRG 2
Mem2 42.4%
Mem CD161 34.5%
Mem1 16.7%
TRG 4
Mem2 27.7%
Mem CD161 29.8%
Mem1 29.1%
CD69
CD161
c
CD4 Mem2
Frequency of CD4 Memory T cells (%)
**
R= -0.646
TRG
Pathological response
d
Mem1
Mem2
Mem CD161
Frequency of CD4 (%)
PN IM TC

**Extended Data Fig. 6: Manual gating confirms the association of CD4 Mem2 with response to neoadjuvant treatment.** (a) Gating strategy for CD4 memory T cells (live single cells $CD45^+$ $CD3^+$ $CD4^+$ $CD45RO^+$ $CD127^+$ $FOXP3^-$ $CD103^-$ $CD38^-$) separating CD4 Mem1 ($CD69^-$ $CD161^-$), CD4 Mem2 ($CD69^+$ $CD161^-$) and CD4 Mem CD161 ($CD69^+$ $CD161^+$). (b) Representative FACS plots showing CD4 memory composition in a responder (TRG 2) and non-responder (TRG 4). (c) Frequency of manually gated CD4 Mem2 cells among total CD4 memory T cells across TRG groups. Statistical significance was assessed using Spearman correlation; correlation coefficient (R) and P values are shown (**P <0.01). (d) Frequency of the CD4 Mem1, Mem2 and Mem CD161 subsets among total CD4 T cell across perilesional normal (PN), invasive margin (IM) and tumor center (TC). Statistical significance was assessed by paired Wilcoxon tests with Benjamini-Hochberg correction. Adjusted P values are shown as *P <0.05, **P <0.01, ***P < 0.001,****P <0.0001.

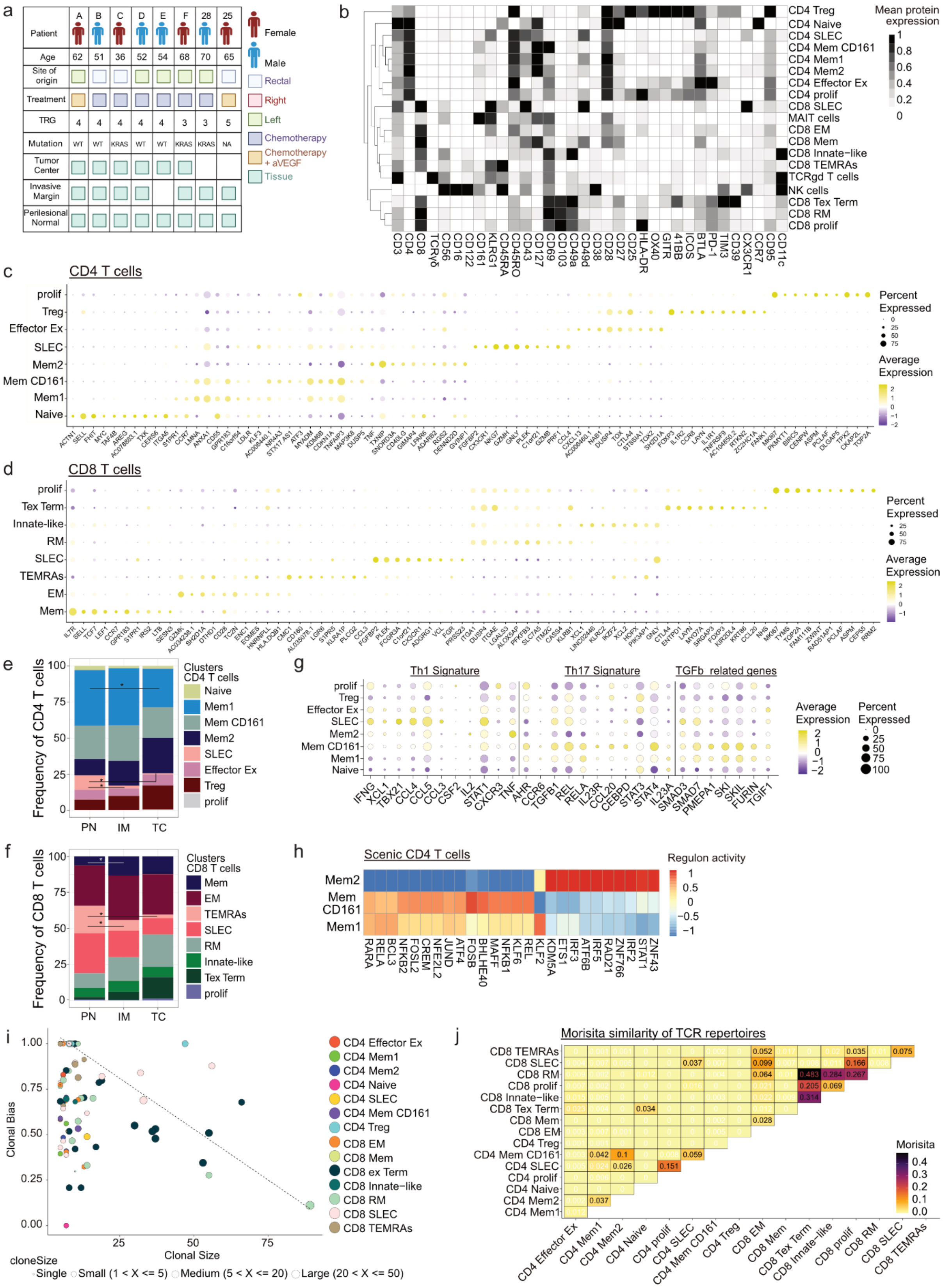

a
Patient
A B C D E F 28 25
Age
62 51 36 52 54 68 70 65
Site of origin
Treatment
TRG
4 4 4 4 4 3 3 5
Mutation
WT WT KRAS WT WT KRAS KRAS NA
Tumor Center
Invasive Margin
Perilesional Normal
Female
Male
Rectal
Right
Left
Chemotherapy
Chemotherapy + aVEGF
Tissue
b
Mean protein expression
CD4 Treg
CD4 Naive
CD4 SLEC
CD4 Mem CD161
CD4 Mem1
CD4 Mem2
CD4 Effector Ex
CD4 prolif
CD8 SLEC
MAIT cells
CD8 EM
CD8 Mem
CD8 Innate−like
CD8 TEMRAs
TCRgd T cells
NK cells
CD8 Tex Term
CD8 RM
CD8 prolif
c
CD4 T cells
prolif
Treg
Effector Ex
SLEC
Mem2
Mem CD161
Mem1
Naive
Percent Expressed
Average Expression
d
CD8 T cells
prolif
Tex Term
Innate-like
RM
SLEC
TEMRAs
EM
Mem
Percent Expressed
Average Expression
e
Frequency of CD4 T cells
Clusters CD4 T cells
Naive
Mem1
Mem CD161
Mem2
SLEC
Effector Ex
Treg
prolif
PN IM TC
g
Th1 Signature
Th17 Signature
TGFb related genes
Average Expression
Percent Expressed
f
Frequency of CD8 T cells
Clusters CD8 T cells
Mem
EM
TEMRAs
SLEC
RM
Innate-like
Tex Term
prolif
PN IM TC
h
Scenic CD4 T cells
Regulon activity
Mem2
Mem CD161
Mem1
i
Clonal Bias
Clonal Size
cloneSize
Single
Small (1 < X <= 5)
Medium (5 < X <= 20)
Large (20 < X <= 50)
CD4 Effector Ex
CD4 Mem1
CD4 Mem2
CD4 Naive
CD4 SLEC
CD4 Mem CD161
CD4 Treg
CD8 EM
CD8 Mem
CD8 ex Term
CD8 Innate−like
CD8 RM
CD8 SLEC
CD8 TEMRAs
j
Morisita similarity of TCR repertoires
Morisita

**Extended Data Fig. 7: Combined transcriptomic and proteomic profiling highlights a Th1 helper signature and CD8 progenitor-like features associated with response to neoadjuvant treatment.** (a) Clinical information available across 21 samples from eight patients included in the Rhaspody analysis. (b) Heatmap showing mean normalized protein expression per cluster across 19 lymphocyte populations. (c, d) DotPlot showing mean expression and proportion of cell expressing the top 10 differentially expressed genes (log fold change > 0.25, detection rate ≥ 25%) for CD4 (c) and CD8 (d) T cell clusters. (e, f) Bar plots showing mean frequencies of CD4 (e) and CD8 (f) T cell subsets across patients in each region, normalized so that subsets sum to 100% per region. Statistical significance was assessed using paired Wilcoxon tests with Benjamini-Hochberg correction (*P value adjusted <0.05). (g) Dot Plot showing expression of genes associated with Th1, Th17 and TGF-β related signature. (h) Heatmap showing scaled regulon activity of the 10 most significant regulons across CD4 Mem1, CD4 Mem CD161 and CD4 Mem2 subsets (Wilcoxon rank-sum test with Benjamini-Hochberg correction) accessed using SCENIC. (i) Clonal bias analysis showing distribution of individual TCR clones across cellular compartments and clonal sizes per patient. (J) Morisita index measuring strict TCR sharing between CD4 and CD8 subsets.

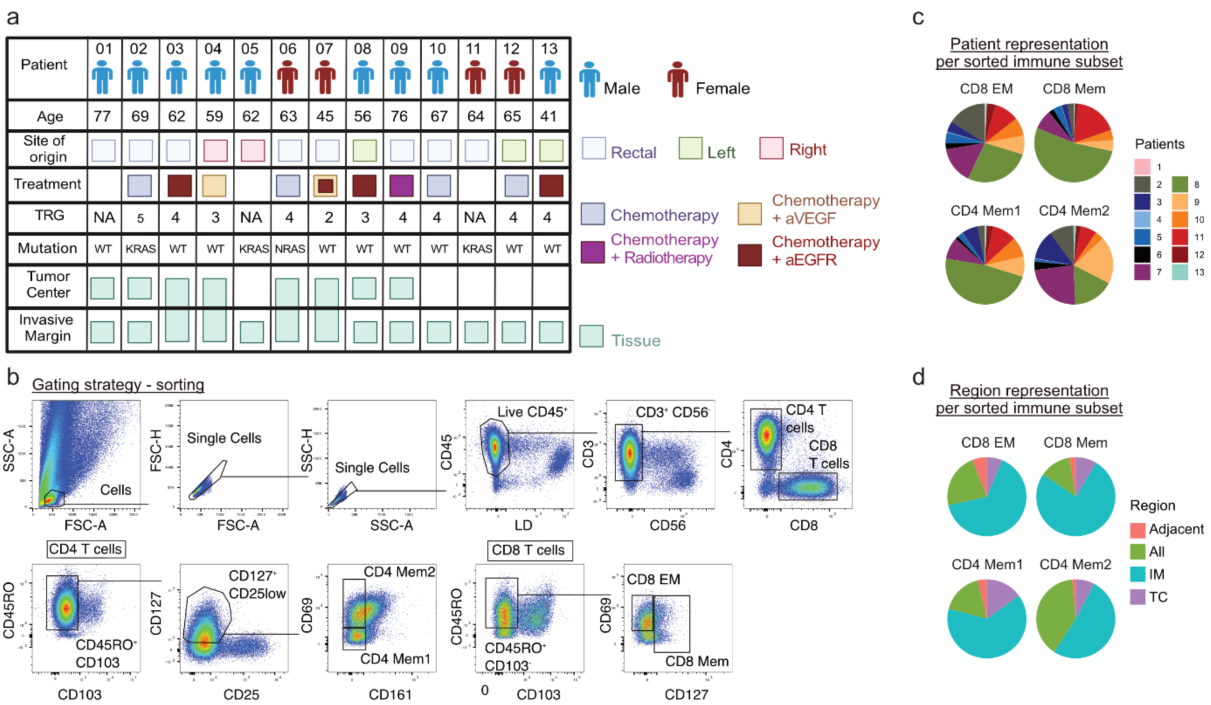
a
Patient
01
02
03
04
05
06
07
08
09
10
11
12
13
Male
Female
Age
77
69
62
59
62
63
45
56
76
67
64
65
41
Site of origin
Rectal
Left
Right
Treatment
Chemotherapy
Chemotherapy + aVEGF
Chemotherapy + Radiotherapy
Chemotherapy + aEGFR
TRG
NA
5
4
3
NA
4
2
3
4
4
NA
4
4
Mutation
WT
KRAS
WT
WT
KRAS
NRAS
WT
WT
WT
WT
KRAS
WT
WT
Tumor Center
Invasive Margin
Tissue
b
Gating strategy - sorting
SSC-A
FSC-A
Cells
FSC-H
Single Cells
SSC-H
SSC-A
Single Cells
CD45
LD
Live CD45+
CD3
CD56
CD3+ CD56-
CD4
CD8
CD4 T cells
CD8 T cells
CD4 T cells
CD45RO
CD103
CD45RO+ CD103-
CD127
CD25
CD127+ CD25low
CD69
CD161
CD4 Mem2
CD4 Mem1
CD8 T cells
CD45RO
CD103
CD45RO+ CD103-
0
CD69
CD127
CD8 EM
CD8 Mem
c
Patient representation per sorted immune subset
CD8 EM
CD8 Mem
CD4 Mem1
CD4 Mem2
Patients
1
2
3
4
5
6
7
8
9
10
11
12
13
d
Region representation per sorted immune subset
CD8 EM
CD8 Mem
CD4 Mem1
CD4 Mem2
Region
Adjacent
All
IM
TC

**Extended Data Fig. 8: CD4 Mem2 T cells exhibit superior Th1 activation and cytokine production, while CD8 Mem and EM cells show complementary functional roles.** (a) Clinical information from 13 patients included in the *ex vivo* stimulation analysis. (b) Gating strategy for CD4 memory cells (live single cells $CD45^+$ $CD3^+$ $CD4^+$ $CD45RO^+$ $CD127^+$ $CD25^-$ $CD103^-$) separating CD4 Mem1 ($CD69^-$ $CD161^-$), CD4 Mem2 ($CD69^+$ $CD161^-$), CD8 Mem ($CD45RO^+$ $CD127^+$) and CD8 EM ($CD45RO^+$ $CD69^+$ $CD127^-$). (c) Pie chart showing pooled immune subset composition by patient. (d) Pie chart showing pooled immune subset composition by region.

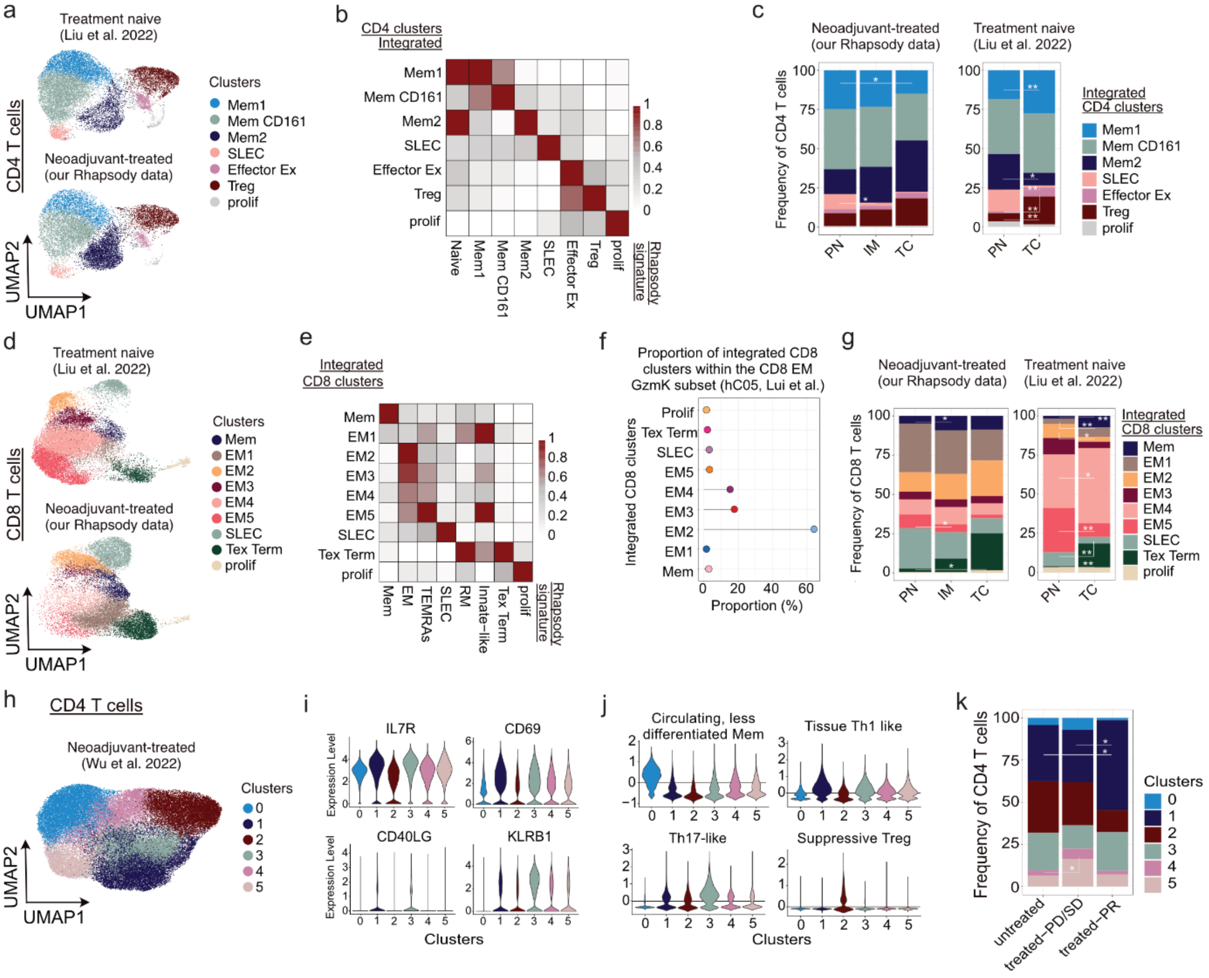
a
Treatment naive
(Liu et al. 2022)
Neoadjuvant-treated
(our Rhapsody data)
CD4 T cells
UMAP2
UMAP1
Clusters
Mem1
Mem CD161
Mem2
SLEC
Effector Ex
Treg
prolif
b
CD4 clusters
Integrated
Rhapsody signature
Naive
Mem1
Mem CD161
Mem2
SLEC
Effector Ex
Treg
prolif
c
Neoadjuvant-treated
(our Rhapsody data)
Treatment naive
(Liu et al. 2022)
Frequency of CD4 T cells
PN
IM
TC
Integrated
CD4 clusters
d
CD8 T cells
e
Integrated
CD8 clusters
Mem
EM1
EM2
EM3
EM4
EM5
SLEC
Tex Term
prolif
EM
TEMRAs
RM
Innate−like
f
Proportion of integrated CD8 clusters within the CD8 EM GzmK subset (hC05, Lui et al.)
Integrated CD8 clusters
Proportion (%)
g
Frequency of CD8 T cells
Integrated
CD8 clusters
h
CD4 T cells
Neoadjuvant-treated
(Wu et al. 2022)
i
IL7R
CD69
CD40LG
KLRB1
Expression Level
Clusters
j
Circulating, less differentiated Mem
Tissue Th1 like
Th17-like
Suppressive Treg
k
Frequency of CD4 T cells
untreated
treated−PD/SD
treated−PR

**Extended Data Fig. 9: Single-cell integration reveals conserved and therapy-responsive T cell states in CRLM.** (a) UMAPs of CD4 T cells from untreated or neoadjuvant-treated CRLM patients, integrating this study (15,470 cells) with Liu et al., 2022 (17,967 cells), showing seven clusters separated by data set. (b) Heatmap of Rhapsody signature scores for CD4 T cell subsets across the seven clusters from the untreated data set. (c) Bar plots showing mean frequencies of CD4 subsets per patients and region, separated by dataset, normalized to 100% per region. (d) UMAPs of CD8 T cells from untreated or neoadjuvant-treated CRLM patients, integrating this study (19,767 cells) with Liu et al., 2022 (22,446 cells), showing nine clusters separated by data set. (e) Heatmap of Rhapsody signature scores for CD8 subsets across the nine clusters from the untreated data set. (f) Lollipop graph showing the proportion of integrated CD8 clusters corresponding to the CD8 EM GzmK subset (hC05 defined in Liu et al., 2022) (g) Bar plots showing mean frequencies of CD8 subsets per patient and region, separated by dataset, normalized to 100% per region. (h) UMAP of CD4 T cells from untreated or neoadjuvant-treated CRLM patients (Wu et al., 2022), showing five clusters. (i) Violin plots showing expression of IL7R, *CD69*, *CD40LG* and *KLRB1* across the five CD4 clusters. (j) Violin plots showing signature scores for circulating, less-differentiated memory, tissue Th1-like, Th17-like, and suppressive Treg programs across five CD4 T cell clusters. (k) Bar plots showing mean frequencies of CD4 T cell subsets across patients stratified as untreated, treated with progressive or stable disease (treated-PD/SD), or treated with partial response (treated-PR), normalized to 100% per region. (c, g, k) Statistical significance was assessed using paired Wilcoxon tests with Benjamini-Hochberg correction. Adjusted P values are show as *P <0.05 and **P <0.01.

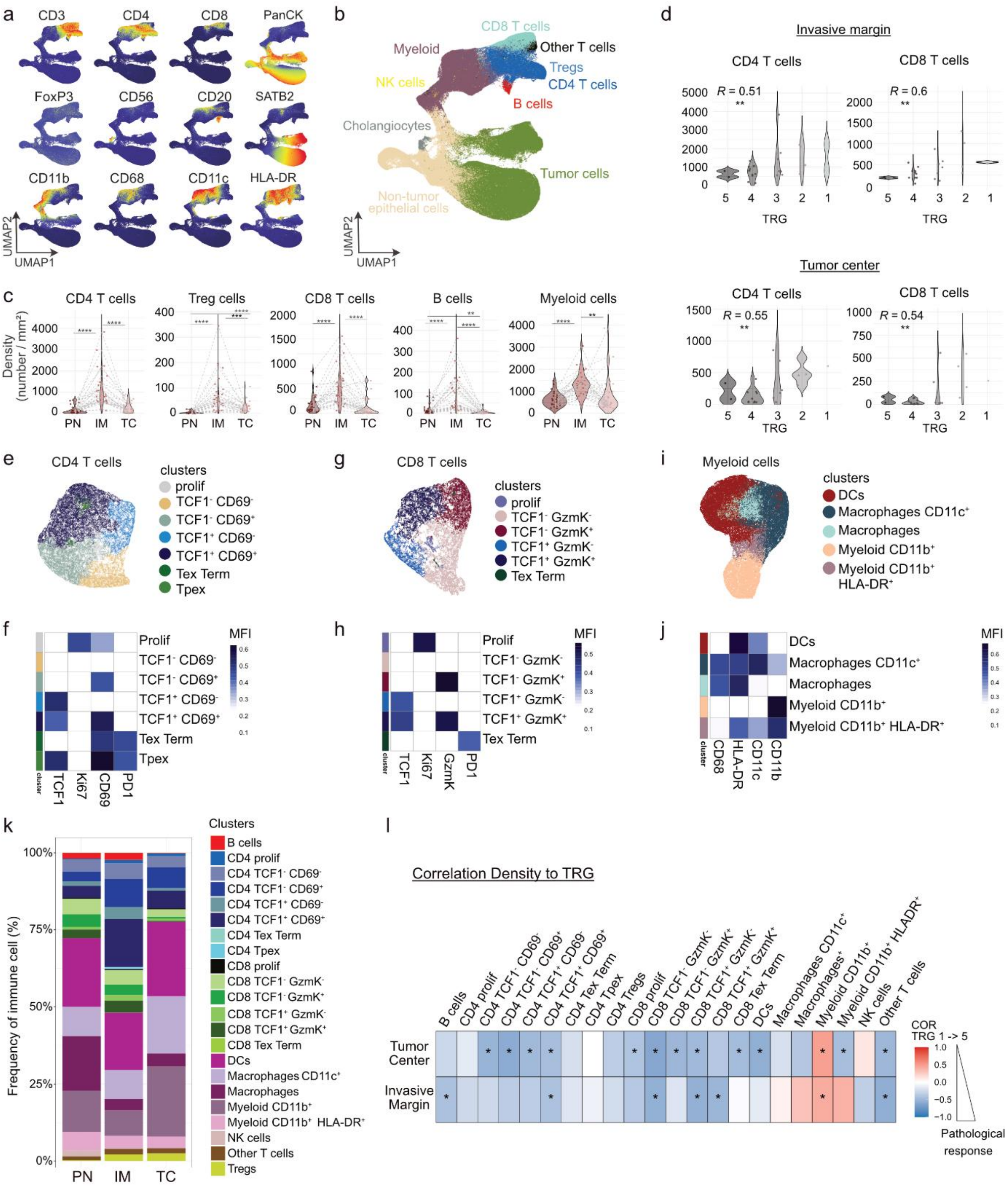

a
CD3
CD4
CD8
PanCK
FoxP3
CD56
CD20
SATB2
CD11b
CD68
CD11c
HLA-DR
UMAP2
UMAP1
b
CD8 T cells
Myeloid
Other T cells
Tregs
NK cells
CD4 T cells
B cells
Cholangiocytes
Tumor cells
Non-tumor epithelial cells
UMAP2
UMAP1
c
CD4 T cells
Treg cells
CD8 T cells
B cells
Myeloid cells
Density (number / mm²)
PN IM TC
d
Invasive margin
CD4 T cells
CD8 T cells
R = 0.51
R = 0.6
TRG
Tumor center
R = 0.55
R = 0.54
e
CD4 T cells
clusters
prolif
TCF1⁻ CD69⁻
TCF1⁻ CD69⁺
TCF1⁺ CD69⁻
TCF1⁺ CD69⁺
Tex Term
Tpex
g
CD8 T cells
clusters
prolif
TCF1⁻ GzmK⁻
TCF1⁻ GzmK⁺
TCF1⁺ GzmK⁻
TCF1⁺ GzmK⁺
Tex Term
i
Myeloid cells
clusters
DCs
Macrophages CD11c⁺
Macrophages
Myeloid CD11b⁺
Myeloid CD11b⁺ HLA-DR⁺
f
MFI
Prolif
TCF1⁻ CD69⁻
TCF1⁻ CD69⁺
TCF1⁺ CD69⁻
TCF1⁺ CD69⁺
Tex Term
Tpex
cluster
TCF1
Ki67
CD69
PD1
h
MFI
Prolif
TCF1⁻ GzmK⁻
TCF1⁻ GzmK⁺
TCF1⁺ GzmK⁻
TCF1⁺ GzmK⁺
Tex Term
cluster
TCF1
Ki67
GzmK
PD1
j
MFI
DCs
Macrophages CD11c⁺
Macrophages
Myeloid CD11b⁺
Myeloid CD11b⁺ HLA-DR⁺
cluster
CD68
HLA-DR
CD11c
CD11b
k
Clusters
B cells
CD4 prolif
CD4 TCF1⁻ CD69⁻
CD4 TCF1⁻ CD69⁺
CD4 TCF1⁺ CD69⁻
CD4 TCF1⁺ CD69⁺
CD4 Tex Term
CD4 Tpex
CD8 prolif
CD8 TCF1⁻ GzmK⁻
CD8 TCF1⁻ GzmK⁺
CD8 TCF1⁺ GzmK⁻
CD8 TCF1⁺ GzmK⁺
CD8 Tex Term
DCs
Macrophages CD11c⁺
Macrophages
Myeloid CD11b⁺
Myeloid CD11b⁺ HLA-DR⁺
NK cells
Other T cells
Tregs
Frequency of immune cell (%)
PN IM TC
l
Correlation Density to TRG
Tumor Center
Invasive Margin
COR TRG 1 -> 5
Pathological response

**Extended Data Fig. 10: Spatial coordination of CD4 memory, CD8 T cells and APCs defines neoadjuvant treatment-responsive niches.** (a, b) UMAPs of all stained cells from TMAs (408,311 cells across 30 patients and 3 regions), colored by normalized marker intensity (a) or by major FlowSom clusters (b). (c) Violin plots showing density (cells/mm$^2$) of major immune clusters across perilesional normal (PN), invasive margin (IM) and tumor center (TC). Statistical significance was assessed by paired Wilcoxon tests with Benjamini-Hochberg correction. Adjusted P values are shown as *P <0.05, **P <0.01, ***P < 0.001, ****P <0.0001. (d) Violin plots showing CD4 and CD8 T cell density per patient in IM and TC, grouped by pathological response (TRG). Positive correlation indicates association with responders (low TRG). (e) UMAP of CD4 T cells (48,803 cells), colored by detailed cluster. (f) Heatmap showing mean normalized marker expression across CD4 T cell subsets. (g) UMAP of CD8 T cells (22,894 cells), colored by detailed cluster. (h) Heatmap showing mean normalized marker expression across CD8 T cell subsets. (i) UMAP of myeloid cells (95,760 cells), colored by detailed cluster. (j) Heatmap showing mean normalized marker expression across myeloid subsets. (k) Bar plots showing median frequencies of immune subtypes per patient across PN, IM and TC. (l) Heatmap showing Spearman correlation coefficients (COR) between immune cell density per patient and TRG, displayed by region. Positive correlation (red) indicate association with non-responders, negative correlation (blue) with responders. (d, l) Statistical significance was assessed using Spearman correlation; correlation coefficient (R) and *p* value are indicated (* *p* <0.05 and ** *p* <0.01).

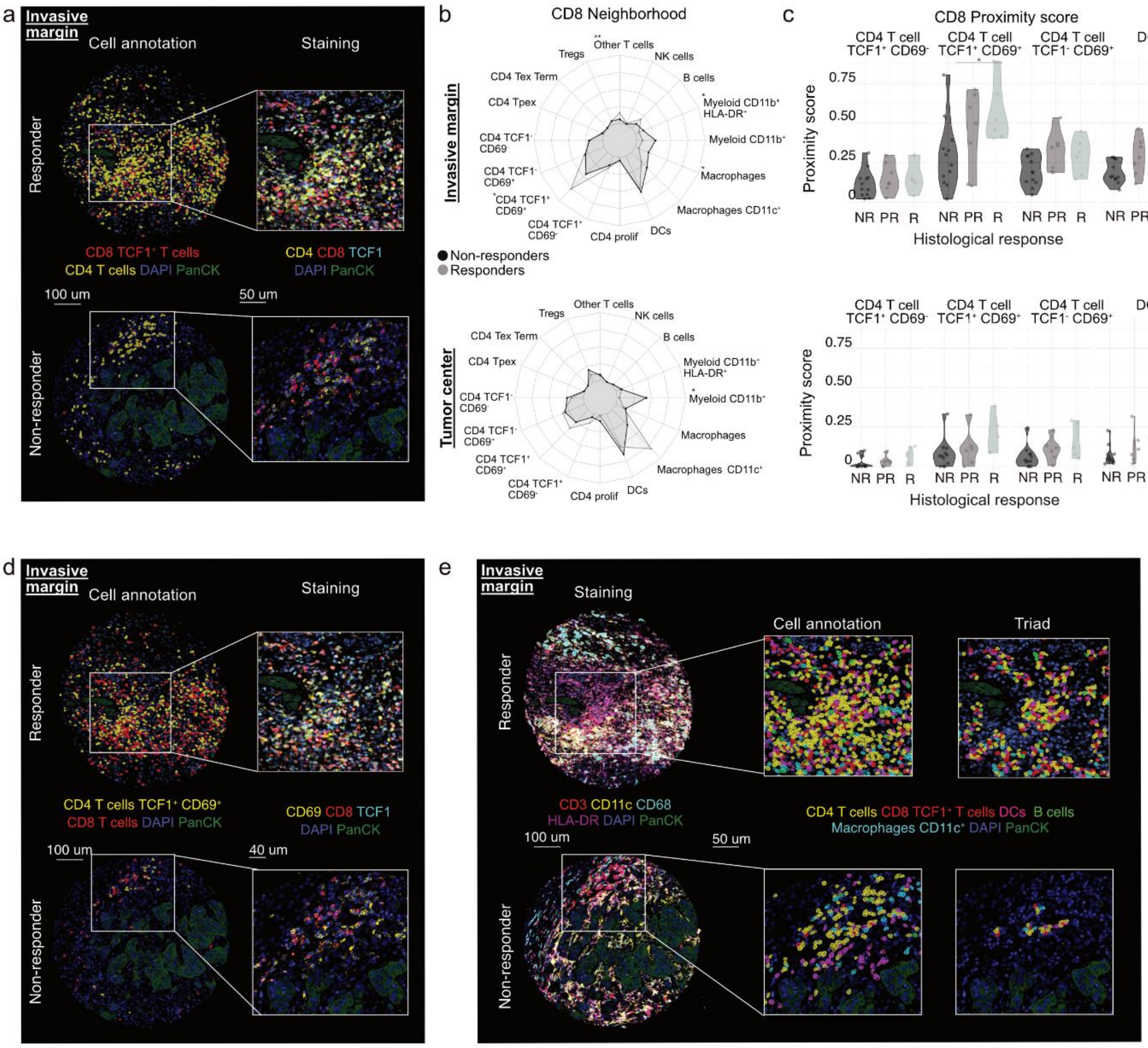

a
Invasive margin
Cell annotation
Staining
Responder
Non-responder
CD8 TCF1+ T cells
CD4 T cells DAPI PanCK
CD4 CD8 TCF1
DAPI PanCK
100 um
50 um
b
CD8 Neighborhood
Invasive margin
Tumor center
Other T cells
NK cells
B cells
Myeloid CD11b+ HLA-DR+
Myeloid CD11b+
Macrophages
Macrophages CD11c+
DCs
CD4 prolif
CD4 TCF1+ CD69-
CD4 TCF1+ CD69+
CD4 TCF1- CD69+
CD4 TCF1- CD69
CD4 Tpex
CD4 Tex Term
Tregs
Non-responders
Responders
c
CD8 Proximity score
CD4 T cell TCF1+ CD69-
CD4 T cell TCF1+ CD69+
CD4 T cell TCF1- CD69+
DCs
Proximity score
0.75
0.50
0.25
0
NR PR R
Histological response
d
Invasive margin
Cell annotation
Staining
Responder
Non-responder
CD4 T cells TCF1+ CD69+
CD8 T cells DAPI PanCK
CD69 CD8 TCF1
DAPI PanCK
100 um
40 um
e
Invasive margin
Staining
Cell annotation
Triad
Responder
Non-responder
CD3 CD11c CD68
HLA-DR DAPI PanCK
CD4 T cells CD8 TCF1+ T cells DCs B cells
Macrophages CD11c+ DAPI PanCK
100 um
50 um

**Extended Data Fig. 11: Spatial coordination of CD4 memory, CD8 T cells and APCs defines neoadjuvant treatment-responsive niches.** (a) Representative images from the invasive margin of a responder (TRG 2) and non-responder (TRG 4). Left: CD4 T cells and CD8 TCF1$^{+}$ GzmK$^{+/-}$ T cells. Right: Zoom-in showing staining of selected markers used for phenotyping. (b) Radar plots showing mean composition of the 10 closest non-CD8 neighbors for each CD8 T cells in invasive margin and tumor center, stratified by response. Radial axes are scaled to a maximum of 35%. Statistical significance was assessed by permutation-based independence tests with Benjamini–Hochberg correction (*adjusted P < 0.05, ** < 0.01). (c) Violin plots showing proximity scores of neighbor cell types to CD8 T cells, per region and response group. Statistical significance was tested separately for each neighbor type and region, by Kruskal–Wallis followed by Benjamini–Hochberg-adjusted Wilcoxon rank-sum tests (*adjusted P <0.05, ** <0.01). (d) Representative images from the invasive margin of a responder (TRG 2) and non-responder (TRG 4). Left: CD8 T cells and CD4 TCF1$^{+}$ CD69$^{+}$ T cells. Right: Zoom-in showing staining of selected markers used for phenotyping. (e) Representative image from the invasive margin of a responder (TRG 2) and non-responder (TRG 4). Left: myeloid and lymphoid marker staining across the TMA, Middle: Zoom-in showing CD4 T cells, CD8 TCF1$^{+}$ GzmK$^{+/-}$ T cells, B cells, Macrophages CD11c$^{+}$ or DCs. Right: Zoom-in showing cells participating in at least one qualifying triad.

**Extended Data Table 1: Antibodies for Flow cytometry**

| Antibody target, fluorochrome and clone | Dilution | SOURCE | IDENTIFIER |
|---|---|---|---|
| CXCR6 BV650 clone 13B 1E5 | 50 | BD Biosciences | 743600 |
| CCR7 BV785 clone G043H7 | 100 | Biolegend | 353230 |
| CX3CR1 BUV661 clone 2A9-1 | 200 | BD Biosciences | 750690 |
| CD45RO BUV615 clone UCHL1 | 200 | BD Biosciences | 751167 |
| CD103 Biotin clone Ber-ACT8 | 200 | Biolegend | 350220 |
| CD3 BUV805 clone UCHT-1 | 75 | BD Biosciences | 612895 |
| CD8 BV570 clone RPA-T8 | 100 | Biolegend | 301038 |
| TCRgd PE-Cy5 clone IMMU510 | 20 | Beckman Coulter | IM2662U |
| CD45 PerCp clone HI-30 | 100 | Biolegend | 304025 |
| CD16 BUV496 clone 3G8 | 100 | BD Biosciences | 612944 |
| CD161 BUV563 clone HP-3G10 | 100 | BD Biosciences | 749223 |
| CD19 Super Bright 436 clone HIB19 | 50 | Thermo Fisher Scientific | 62-0199-42 |
| CD56 PE/Dazzle 594 clone 5.1H11 | 200 | Biolegend | 362543 |
| CD4 Spark Blue 550 clone SK3 | 200 | Biolegend | 344656 |
| CD14 Spark NIR clone 63D3 | 200 | Biolegend | 367150 |
| PD-1 BV605 clone EH12.2H7 | 40 | Biolegend | 329924 |
| CD38 APC-Fire 810 clone HIT2 | 100 | Biolegend | 303550 |
| CD69 BV421 clone FN50 | 100 | Biolegend | 310930 |
| ICOS BV750 clone C398.4A | 200 | Biolegend | 313558 |
| TIGIT BUV395 clone 741182 | 50 | BD Biosciences | 747845 |
| KLRG1 PE-Fire 810 clone SA231A2 | 400 | Biolegend | 367733 |
| TIM3 BB700 clone 344823 | 200 | BD Biosciences | 747957 |
| CD39 BUV737 clone TU66 | 100 | BD Biosciences | 564726 |
| CD127 BV510 clone A019D5 | 75 | Biolegend | 351332 |
| CTLA-4 BB790 clone BNI3 | 100 | BD Biosciences | 624296 |
| EOMES APC-eFluor 780 clone WD1928 | 40 | Thermo Fisher Scientific | 47-4877-41 |
| T-bet BV711 clone 4B10 | 50 | Biolegend | 644820 |
| Ki67 BV480 clone SolA15 | 200 | Thermo Fisher Scientific | 414-5698-82 |
| FoxP3 PE-Cy7 clone 236A/E7 | 25 | Thermo Fisher Scientific | 25-4777-42 |
| TOX PE clone REA473 | 200 | Miltenyi Biotec | 130-120-716 |
| Granzyme B AlexaFluor 700 clone GB11 | 250 | BD Biosciences | 560213 |
| TCF-1 AlexaFluor 488 clone C63D9 | 300 | Cell Signaling | 6444S |
| Streptavidin BB630-P2 | 500 | BD Biosciences | 624294 |

**Extended Data Table 2: Antibodies for Rhapsody**

| AbSeq Oligo | Dilution | SOURCE | IDENTIFIER |
|---|---|---|---|
| AbSeq Immune Discovery Panel | | BD Biosciences | 625970 |
| Hu CD38 Ab-O AHS0022 HIT2 | 100 | BD Biosciences | 940057 |
| Hu CD49d Ab-O AHS0063 9F10 | 100 | BD Biosciences | 940059 |
| Hu CD103 Ab-O AHS0001 BER-ACT8 | 100 | BD Biosciences | 940067 |
| Hu CD49a Ab-O AHS0101 SR84 | 100 | BD Biosciences | 940094 |
| Hu CX3CR1 Olgo AHS0125 2A9-1 | 100 | BD Biosciences | 940216 |
| Hu CD122 Olgo AHS0146 MIK-BETA3 | 100 | BD Biosciences | 940232 |
| Hu CD43 Olgo AHS0200 1G10 | 100 | BD Biosciences | 940278 |
| Hu CD95 Ab-O AHS0023 DX2 | 100 | BD Biosciences | 940037 |
| Hu CD45RO Ab-O AHS0036 UCHL1 | 100 | BD Biosciences | 940022 |
| Hu CD69 Ab-O AHS0010 FN50 | 100 | BD Biosciences | 940019 |
| Hu TCRgd Ab-O AHS0015 B1 | 100 | BD Biosciences | 940057 |
| KLRG1 APC clone SA231A2 | 200 | Biolegend | 367715 |
| HU APC Olgo AHS0295 E30-221 | 100 | BD Biosciences | 460078 |
| Hu CD39 Ab-O AHS0006 TU66 | 100 | BD Biosciences | 940073 |

**Extended Data Table 3: Antibodies for ex vivo stimulation**

| Antibody target, fluorochrome and clone | Dilution | SOURCE | IDENTIFIER |
|---|---|---|---|
| Anti-human CD3 BUV805 clone UCHT-1 | 75 | BD Biosciences | 612895 |
| Anti-human CD69 BV421 clone FN50 | 100 | Biolegend | 310930 |
| Anti-human CD103 BV510 clone Ber-ACT8 | 50 | BD Biosciences | 743651 |
| Anti-human CD127 BV605 clone A019D5 | 50 | Biolegend | 351333 |
| Anti-human CD161 BV785 clone HP-3G10 | 100 | Biolegend | 339930 |
| Anti-human CD4 Spark Blue 550 clone SK3 | 200 | Biolegend | 344656 |
| Anti-human CD45 PerCp clone HI-30 | 100 | Biolegend | 304025 |
| Anti-human CD8 PE clone SK-1 | 100 | BD Biosciences | 345773 |
| Anti-human CD56 PE/Dazzle 594 clone 5.1H11 | 200 | Biolegend | 362543 |
| Anti-human CD45RO PE-Cy7 clone UCHL1 | 200 | Biolegend | 304230 |
| Anti-human CD25 R718 clone BC96 | 50 | BD Biosciences | 567575 |
| Anti-human HLA-DR BUV395 clone G46-6 | 100 | BD Biosciences | 564040 |
| Anti-human CD27 BUV563 clone M-T271 | 150 | BD Biosciences | 741366 |
| Anti-human Ki67 BV480 clone SolA15 | 200 | Thermo Fisher Scientific | 414-5698-82 |
| Anti-human PD-1 BV605 clone EH12.2H7 | 40 | Biolegend | 329924 |
| Anti-human T-bet BV711 clone 4B10 | 50 | Biolegend | 644820 |
| Anti-human CD28 BV750 clone CD28.2 | 100 | BD Biosciences | 747329 |
| Anti-human TIM3 VioBright FITC clone F38.2E | 50 | Miltenyi | 130-104-687 |
| Anti-human CD25 PE clone M-A251 | 100 | Biolegend | 356103 |
| Anti-human Granzyme K PE-Cy7 clone GM26E7 | 100 | Biolegend | 370515 |
| Anti-human CD40L AF647 clone 24-31 | 100 | Biolegend | 310818 |
| Anti-human Granzyme B AF700 clone GB11 | 300 | BD Biosciences | 560213 |
| Anti-human CD38 APC-Fire 810 clone HIT2 | 150 | Biolegend | 303550 |

**Extended Data Table 4: Antibodies for multiplex imaging**

| Antibody target, fluorochrome and clone | SOURCE | IDENTIFIER |
|---|---|---|
| Anti-human CD3 rabbit | Lunaphore | MR10010 |
| Anti-human FoxP3 mouse | Lunaphore | MR10040 |
| Anti-human CD4 rabbit | Lunaphore | MR10020 |
| Anti-human HLA-DR mouse | DAKO | M0746 |
| Anti-human CD56 rabbit | Lunaphore | MR10060 |
| Anti-human CD8 mouse | DAKO | M7103 |
| Anti-human CD11c rabbit | Lunaphore | MR10070 |
| Anti-human CD68 mouse | Invitrogen | MA5-13324 |
| Anti-human CD20 mouse | Lunaphore | L26 |
| Anti-human PD-1 rabbit | Lunaphore | MR10120 |
| Anti-human aSMA mouse | DAKO | M0851 |
| Anti-human STAB2 rabbit | Diagnostic Biosystem | RMAB112 |
| Anti-human Granzyme B rabbit | Abcam | ab208586 |
| Anti-human Pan Cytokeratin mouse | Lunaphore | MR10140 |
| Anti-human CD69 rabbit | Abcam | 310902 |
| Anti-human TCF1 rabbit | Cell Signaling | 2203 |
| Anti-human Granzyme K rabbit | Abcam | ab282703 |
| Anti-human CD11b rabbit | Cell Signaling | 48893 |
| Anti-human Ki67 rabbit | Abcam | ab224639 |

**Extended Data Table 5. Cofactor used in Flow Cytometry analysis for marker expression transformation.**

| Marker | cofactor used |
|---|---|
| CD38 | 2500 |
| Eomes | 16000 |
| TCF1 | 15000 |
| GzmB | 1700 |
| TCRVa7.2 | 3500 |
| CD103 | 2300 |
| TIM3 | 2700 |
| CTLA4 | 3500 |
| TIGIT | 1300 |
| CD16 | 2 |
| CD161 | 2000 |
| CD45RO | 900 |
| CX3CR1 | 1300 |
| CD39 | 1300 |
| CD3 | 1500 |
| CD69 | 1700 |
| Ki67 | 2 |

| | |
|---|---|
| CD127 | 2800 |
| CD8 | 2 |
| PD1 | 50 |
| CXCR6 | 1600 |
| Tbet | 10000 |
| ICOS | 2500 |
| CCR7 | 7000 |
| TOX | 30000 |
| TCRgd | 2500 |
| CD45RA | 4500 |
| FoxP3 | 8500 |
| CD56 | 2500 |
| KLRG1 | 2700 |
| CD45 | 4000 |
| CD4 | 2500 |
| CD14 | 4000 |
| CD19 | 2500 |

**Extended Data Table 9. Cofactor used in Multi-plex analysis for marker expression transformation.**

Before cleaning

| marker | cofactor |
| --- | --- |
| FOXP3_nucleus | 40 |
| SATB2_nucleus | 10 |
| GranzymeB_nucleus | 10 |
| TCF1_nucleus | 10 |
| Ki67_nucleus | 10 |
| CD3_cell | 40 |
| HLA-DR_cell | 100 |
| CD4_cell | 50 |
| CD56_cell | 50 |
| CD8_cell | 10 |
| CD11c_cell | 100 |
| CD68_nucleus | 75 |
| PanCK_cell | 25 |
| PD1_cell | 10 |
| CD69_cell | 25 |
| CD20_cell | 10 |
| GranzymeK_nucleus | 10 |
| aSMA_cell | 25 |
| CD11b_cell | 10 |

After cleaning on immune cells

| marker | cofactor |
| --- | --- |
| FOXP3_nucleus | 200 |
| SATB2_nucleus | 10 |
| GranzymeB_nucleus | 10 |
| TCF1_nucleus | 10 |
| Ki67_nucleus | 10 |
| CD3_cell | 40 |
| HLA-DR_cell | 120 |
| CD4_cell | 100 |
| CD56_cell | 100 |
| CD8_cell | 10 |
| CD11c_cell | 200 |
| CD68_nucleus | 50 |
| PanCK_cell | 75 |
| PD1_cell | 100 |
| CD69_cell | 50 |

| | |
|---|---|
| CD20_cell | 50 |
| GranzymeK_nucleus | 10 |
| aSMA_cell | 25 |
| CD11b_cell | 10 |
